# High-affinity detection of endogenously biotinylated neuroligin-1 at excitatory and inhibitory synapses using a tagged knock-in mouse strain

**DOI:** 10.1101/2024.06.11.598408

**Authors:** Charles Ducrot, Adèle Drouet, Béatrice Tessier, Chloé Desquines, Rania-Cérine Mazzouzi, Alexandre Favereaux, Mathieu Letellier, Olivier Thoumine

## Abstract

Neuroligins (NLGNs) are important cell adhesion molecules mediating trans-synaptic contacts between neurons. However, the high-yield biochemical isolation and visualization of endogenous NLGNs have been hampered by the lack of efficient antibodies to these proteins. Thus, to reveal their sub-cellular distribution, binding partners, and synaptic function, NLGNs have been extensively manipulated using knock-down, knock-out, or over-expression approaches, overall leading to controversial results. As an alternative to the manipulation of NLGN expression level, we describe here the generation of a new transgenic mouse strain in which native NLGN1 was N-terminally tagged with a small biotin acceptor peptide (bAP) that can be enzymatically biotinylated by the exogenous delivery of biotin ligase. After showing that knock-in mice exhibit normal behavior as well as similar synaptic number, ultrastructure, transmission properties, and protein expression levels when compared to wild type counterparts, we exploited the fact that biotinylated bAP-NLGN1 can be selectively isolated or visualized using high-affinity streptavidin conjugates. Using immunoblotting and immunofluorescence, we show that bAP-NLGN1 binds both PSD-95 and gephyrin and distributes equally well at excitatory and inhibitory synapses, challenging the historical view that NLGN1 is exclusively localized at excitatory synapses. Using super-resolution fluorescence microscopy and electron microscopy, we further highlight that bAP-NLGN1 forms in the synaptic cleft a subset of nanodomains each containing a few NLGN1 dimers, while the number of nanodomains per synapse positively scales with the post-synapse size. Overall, our study not only provides a novel, extensively characterized transgenic mouse model which will be made available to the scientific community, but also an unprecedented view of the nanoscale organization of endogenous NLGN1.

## Introduction

The assembly of the trillions of synapses formed between neurons during brain development is controlled by many factors including cell adhesion molecules. Among them, neuroligins (NLGN1-4) play critical roles in synapse formation, and are the target of deleterious genetic mutations in autism spectrum disorders (1, 2). NLGNs form constitutive dimers (3, 4) that interact extracellularly with neurexins (NRXNs) in an isoform- and splice variant-dependent manner (5, 6), as well as with MDGAs which compete with trans-synaptic NRXN-NLGN adhesion (7–10). Intracellularly, NLGNs bind PDZ domain containing scaffolding proteins of excitatory synapses through their C-terminus (11–14), the inhibitory synaptic scaffolding molecule gephyrin through a proximal consensus motif (15, 16), the actin-regulating WAVE complex through a WIRS motif located upstream the C-terminus (17, 18), and a still unidentified regulatory partner through a non-canonical motif located close to the transmembrane domain (19). The synaptic function of NLGNs is also regulated by a number of phosphorylation sites recognized by various serine/threonine and tyrosine kinases (20–24).

Despite the advances mentioned above, the high-yield biochemical isolation and super-resolution imaging of endogenous NLGNs have been hampered by the general lack of efficient antibodies to these proteins. Hence, alternative strategies were developed based on the over-expression or rescue of tagged NLGNs in cultured neurons to either isolate or visualize NLGNs. For example, the expression of HRP-tagged NLGN2 served to characterize the inhibitory synapse proteome (25), while GFP or split-GFP tagged versions of NLGNs were used to localize these proteins and study their dynamics at synapses (26–30). However, the introduction of such relatively large tags can bias the function of a protein by altering interactions with specific partners, and over-expression might bias its localization. Indeed, although endogenous NLGN1 is traditionally associated to excitatory synapses and NLGN2 to inhibitory synapses (31, 32), the over-expression of either NLGN isoform increases the number of both excitatory and inhibitory synapses, associated to a gain of function in synaptic transmission (22, 30, 33). Along this line, the synaptic cleft proteome obtained by over-expressed HRP-NLGN1 reveals both excitatory and inhibitory synaptic interactors (25). Conversely, NLGN1 knock-down decreases both excitatory and inhibitory synaptic puncta (34, 35), overall suggesting that NLGN1 might also be implicated at inhibitory synapses. A decade ago, based on the expression of NLGN1 tyrosine point mutants, we proposed that the presence of NLGN1 at excitatory versus inhibitory synapses might be controlled by the differential binding of NLGN1 to PSD-95 versus gephyrin, regulated by phosphorylation (16, 22).

An alternative to such over-expression and knock-down issues and to the use of large fusion proteins has been to rescue endogenous NLGN1 by recombinant NLGN1 carrying small tags such as the biotin acceptor peptide (bAP), followed by labeling of biotinylated NLGN1 with fluorescent streptavidin to study its membrane dynamics and nanoscale localization (10, 27, 28). However, even when doing so, the knock-down of native proteins with shRNAs or cre-recombinase is often incomplete, and the protein might still be slightly over-expressed. Furthermore, since NLGNs exist as several splice variants, the rescue construct has an exogenous promoter and is only one of the possible splice variants (e.g. NLGN1 carrying both A and B splice sites), such that its specific expression may bias the balance between excitatory and inhibitory synapses (36–38). Therefore, there is a crucial need for efficient approaches to target and manipulate native NLGNs. In this direction, an earlier study reported the generation of a constitutive knock-in mouse strain in which endogenous NLGN1 was fused to an HA tag (39), and that was recently used to visualize NLGN1 at high resolution through expansion microscopy in relation to pre-synaptic NRXNs and neurotransmitteer receptors (40). Although powerful, this approach still suffers from the fact that all brain cells including neurons and astrocytes - which also express NLGNs (41) - are labeled, making the subsequent analysis of individual synapses difficult, and that anti-HA antibodies remain of moderate affinity.

To circumvent these limitations, we describe here the generation of a novel knock-in mouse strain in which endogenous NLGN1 was fused to an N-terminal bAP tag that can be enzymatically biotinylated by the viral delivery or electroporation of ER-resident biotin ligase (BirA^ER^), allowing for targeted detection with high-affinity streptavidin conjugates. We first show that those knock-in mice exhibit similar behavior, brain anatomy, synaptic transmission properties, expression levels of synaptic proteins, as well as synapse number and ultrastructure when compared to wild type counterparts at the same age. We then exploited the biotinylation strategy to show an efficient pull-down of bAP- NLGN1 by streptavidin beads, revealing its binding to both PSD-95 and gephyrin, and the formation of NLGN1/NLGN3 heterodimers. Furthermore, labeling of neurons with fluorescent streptavidin combined with the staining of PSD-95 or gephyrin show that bAP-NLGN1 localizes to both excitatory and inhibitory synapses. Finally, we reveal quantitative details about the nanoscale organization of bAP-NLGN1 at synapses using super-resolution fluorescence imaging as well as electron microscopy.

## Results

### Generation of a novel bAP-NLGN1 knock-in mouse strain

In order to modify endogenous NLGN1 for its subsequent biotinylation, we designed a new transgenic mouse strain in which the *Nlgn1* gene in chromosome 3 was N-terminally flanked by a small linker and a DNA sequence coding for the 15 amino-acid biotin acceptor peptide (bAP) (42), inserted after the signal peptide. This molecular replacement strategy was based on a recombinant construct that we previously validated in rodent cultures (22, 28). The generation of the bAP-NLGN1 knock-in mouse strain (hereafter abbreviated KI) was made at the *Phenomin* mouse clinics in Strasbourg (France) (http://www.phenomin.fr/). The strategy was based on the CRISPR/Cas9 method, using a guide RNA (gRNA) specific for *Nlgn1* **(Fig. 1A and Methods)**. This mouse strain was imported to our animal facility, back-crossed with C57BL/6J mice, and raised to the homozygote genotype. To validate the bAP tag knock-in, we performed a genomic DNA sequencing of *Nlgn1* that displayed the correct inserted sequence **(Fig. S1A, B)**, as well as a genotyping that showed the expected shifts in the PCR products from KI mice as compared to wild type (WT) mice of the same genetic background **(Fig. S1C)**.

**Figure 1.**
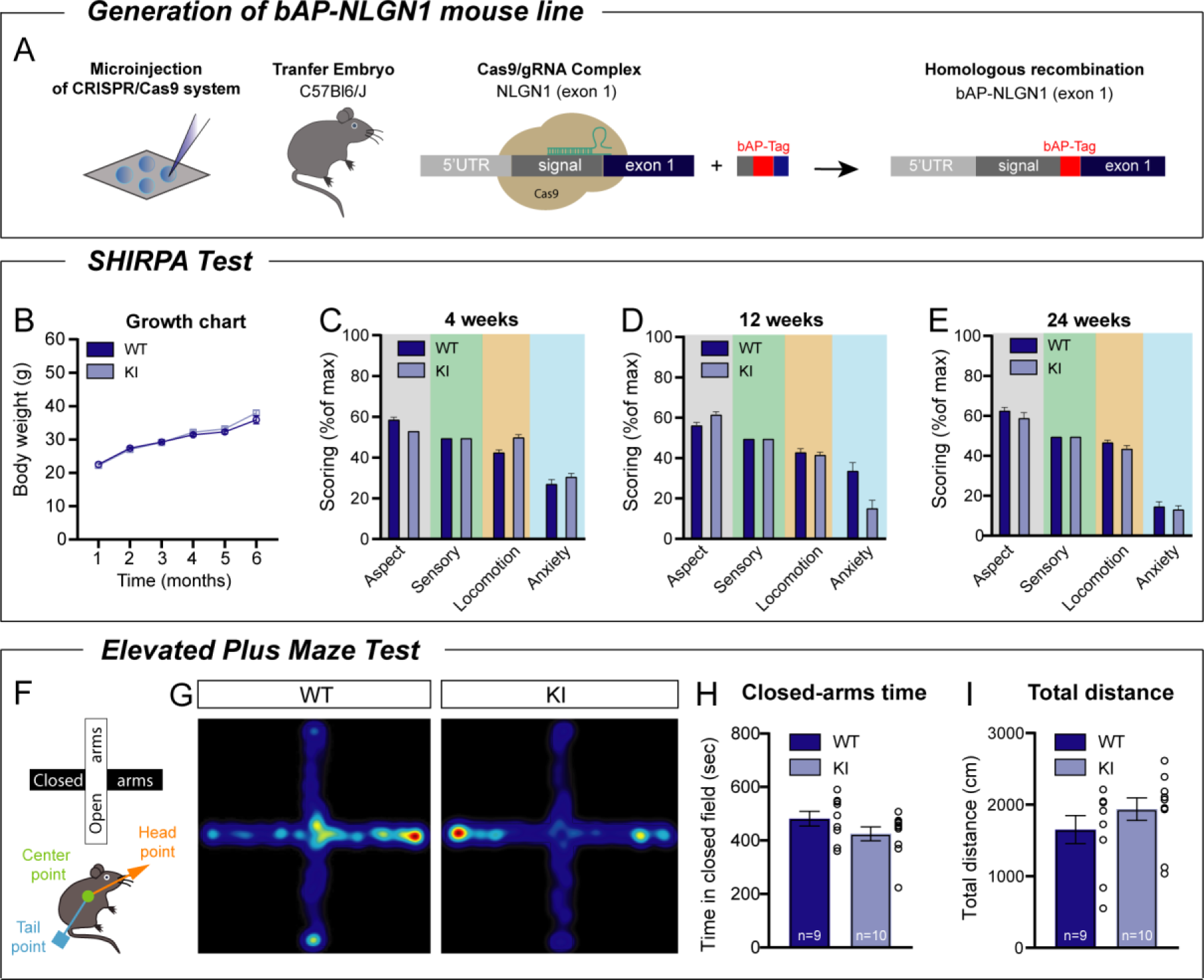
Behavioral characterization of KI mice. **(A)** Schematic representation showing the generation of a novel bAP-NLGN1 knock-in (KI) mouse line by using the CRISPR/Cas9 strategy. **(B)** Growth chart showing the body weight of WT and KI mice over time (1 to 6 months). **(C-E)** Behavioral assessment of WT and KI mice at three different ages (4, 12, and 24 weeks). Grouped analysis was made for each parameter (General aspect, sensory, locomotion and Anxiety) on WT and KI mice. Data were compared by a two-way ANOVA, main effect of genotype, WT versus KI, 4 weeks: *p* = 0.07; 12 weeks: *p* = 0.74 and 24 weeks: *p*=0.33. **(F)** The SHIRPA test was complemented by conducting an elevated plus maze test in adult mice. **(G)** Representative heat maps showing the cumulated positions of a WT and KI mouse, respectively, exploring the elevated plus maze apparatus for 10 min. **(I, J)** The time spend in closed-arms and the total distance travelled, respectively, were quantified and averaged. Data are presented as mean ± SEM. All values were obtained from 9 WT mice and 11 KI mice. Data are presented as mean ± SEM and were compared by Mann-Whitney test (I) *p =* 0.24 and (J) *p =* 0.21. All values were obtained from 9 WT and 10 KI mice.

### Behavioral characterization of KI mice

To check whether the modification of endogenous NLGN1 by insertion of the bAP-tag caused any alteration in mouse physiology or behavior, we quantified a series of parameters by comparing equivalent groups of WT and KI mice bred in the same conditions. First, there was no difference in animal weight between WT and KI groups across a 6-month period **(Fig. 1B)**. We then used the SHIRPA protocol (43) to assess the behavioral phenotype of WT and KI mice at ages of 1 to 6 months, using a wide range of parameters **(Fig. S2)**. Grouped analysis for general aspect, locomotion, and anxiety from 6 months old bAP-NLGN1 mice did not reveal any difference compared to the WT group **(Fig. 1C,E)**. However, a tendency was observed for a decrease in general anxiety parameters of 3-months old KI mice compared to WT mice **(Fig. 1D)**. To specifically monitor anxious behavior, this scoring was complemented by an elevated plus maze test **(Fig. 1F,G)**. No significant difference was observed in the time spend in the closed arms between the WT and KI groups **(Fig 1H)**, nor in the total distance covered by mice during the test **(Fig. 1I)**. Overall, this thorough analysis revealed no major behavioral anomaly of KI mice.

### Normal brain and synapse organization in KI mice

To assess whether the brain of KI mice showed anatomical alterations when compared to WT mice, we performed immunohistochemical stainings of fixed brain sections, focusing on specific regions where NLGN1 is highly expressed (44), i.e. the cortex and hippocampus. We first stained cell nuclei with cresyl violet to visualize cell layers in the cortex **(Fig. 2A,B)**. Analysis of the number of cell nuclei per surface area revealed no difference between sections from WT and KI mice **(Fig. 2C)**. In molecular layers, excitatory and inhibitory pre-synapses were immmunostained for VGluT1 and VGAT **(Fig. 2D,E)**, respectively. The analysis showed no difference in the intensity **(Fig. 2F,G)** or surface area **(Fig. 2H,I)** of the fluorescence signals between sections from WT and KI mice. Similar results were found in hippocampal sections **(Fig. S3)**. To count individual synapses, we cultured primary hippocampal neurons from either WT or KI mice, and immunostained both excitatory synapses with antibodies to VGluT1 and PSD-95 **(Fig. S4A)**, and inhibitory synapses with antibodies to VGAT and gephyrin **(Fig. S4B)**. No difference in the surface density or area of either excitatory or inhibitory synaptic puncta was found between the two genotypes **(Fig. S4C,D)**. Finally, to assess whether synapse ultrastructure is affected in KI mice, we performed transmission electron microscopy (TEM) of ultrathin sections (70 nm) of cortical tissue from either WT or KI mice **(Fig. 2J,K)**. Excitatory and inhibitory synapses of KI mice were not different compared to WT mice in terms of pre-synaptic surface area **(Fig. 2L)**, number of neurotransmitter vesicles **(Fig. 2M)**, number of mitochondria **(Fig. 2N)**, or PSD size **(Fig. 2O)**. Therefore, there was no detectable alteration of brain anatomy, synapse number, or synapse ultrastructure in KI mice.

**Figure 2.**
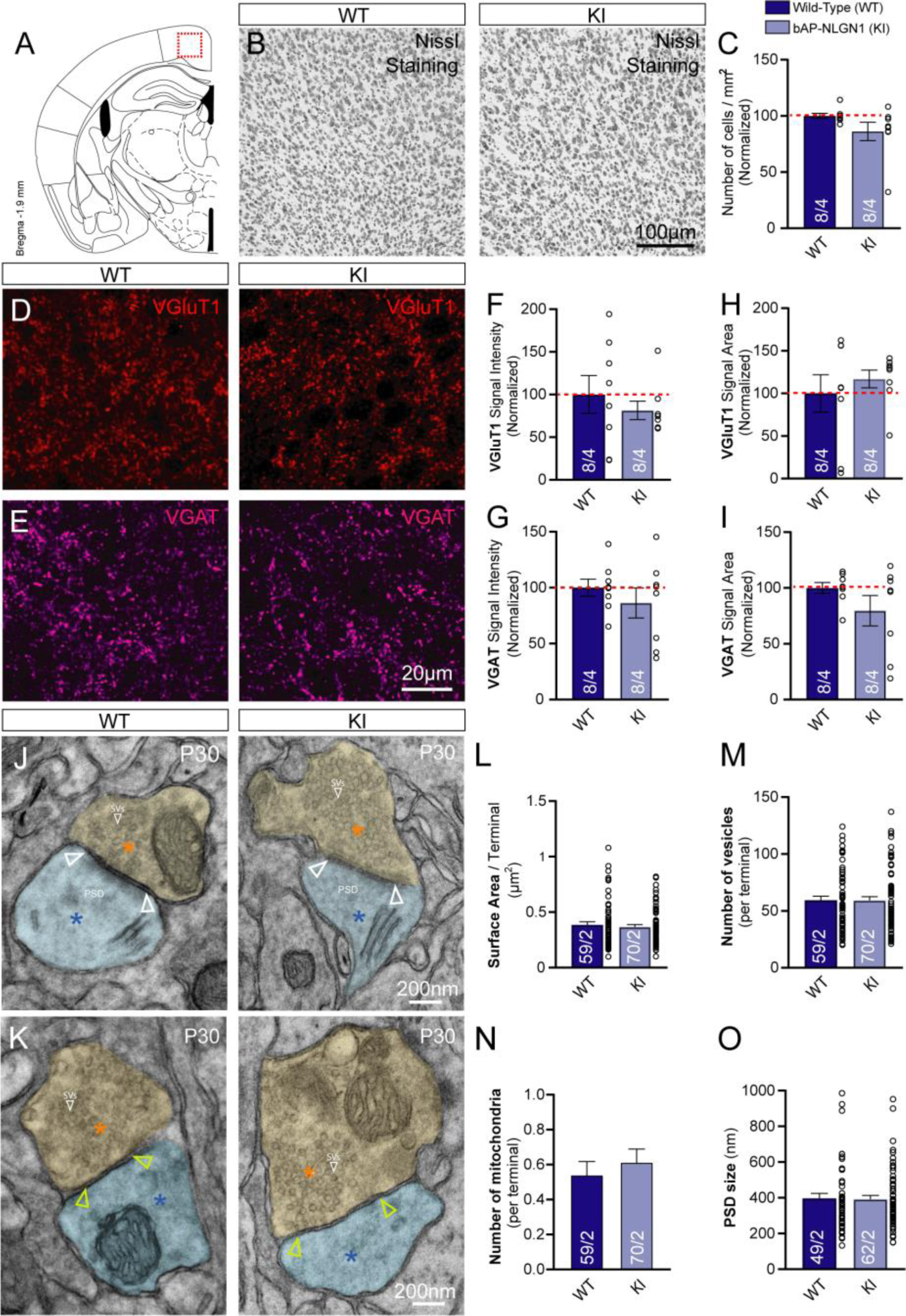
Normal brain development and synapse ultrastructure in KI mice. **((A)** Schematic representation of the cortical slices used for immunohistochemistry characterization. A series of 2 different cortical slices ranging from Bregma -1.5 to -1.9 mm with a total of 4 areas of each hemisphere were analyzed. **(B)** Representative photomicrographs of Nissl staining in cortical brain sections of WT and KI mice. **(C)** Corresponding quantification of the number of cells per mm^2^ in those slices for the two genotypes (*p* = 0.065). **(D, E)** Representative confocal images of immunohistochemical staining for VGLUT1 (red) and VGAT (magenta), respectively, performed on cortical slices from WT and KI mice. Quantification of the signal intensity **(F, G)** and signal surface area **(H, I)**, for VGluT1 and VGAT staining, respectively, normalized to WT levels (in %) (F, *p* = 0.50; G, *p* = 0.64; H, *p* = 0.79 and I, *p* = 0.38). **(J, K)** Representative electron micrographs showing excitatory and inhibitory synapses from WT and KI mice, respectively. The pre- and post-synaptic elements are highlighted in yellow and blue, respectively, and the synaptic cleft is shown with arrows. **(L-N)** Quantification of the surface area (*p*=0.89), number of neurotransmitter vesicles (*p*=0.88), and number of mitochondria of pre-synaptic terminals (*p*=0.59), for both WT and KI mice. **(O)** PSD size of excitatory synapses (*p*=0.84). For Nissl staining and immunolabelling experiments, data represent the mean ± SEM of n = 8 hemispheres from 4 mice for each genotype. For TEM, data represent the mean ± SEM of 59 synapses from 2 WT mice, and 70 synapses from 2 KI mice. In both cases, statistical analyses were carried out by Mann-Whitney test between WT and KI genotypes.

### Normal synaptic transmission in KI mice

To determine whether the bAP tagging of endogenous NLGN1 affected synaptic transmission, we prepared acute hippocampal slices from WT and KI mice and performed patch-clamp recordings of pyramidal CA1 neurons. We stimulated Schaffer’s collaterals originating from CA3 neurons, that form well-characterized synapses onto the dendrites of CA1 neurons (19, 22). By changing the holding membrane potential in voltage-clamp, we recorded sequentially AMPA- and NMDA-receptor mediated EPSCs in response to increasing electrical stimuli **(Fig. 3A)**. There was no difference of evoked EPSCs **(Fig. 3B,C)**, NMDA/AMPA ratio **(Fig. 3D)**, or paired pulse ratio (PPR) **(Fig. 3E,F)** between neurons from WT or KI mice. We also measured spontaneous AMPA receptor-mediated EPSCs occurring in sweeps between stimulations, and found no difference in either the amplitude or frequency of those events between neurons from WT or KI mice **(Fig. 3G-I)**. Thus, excitatory synaptic transmission is not altered in KI mice, at least at the level of this canonical synapse in the hippocampus.

**Figure 3.**
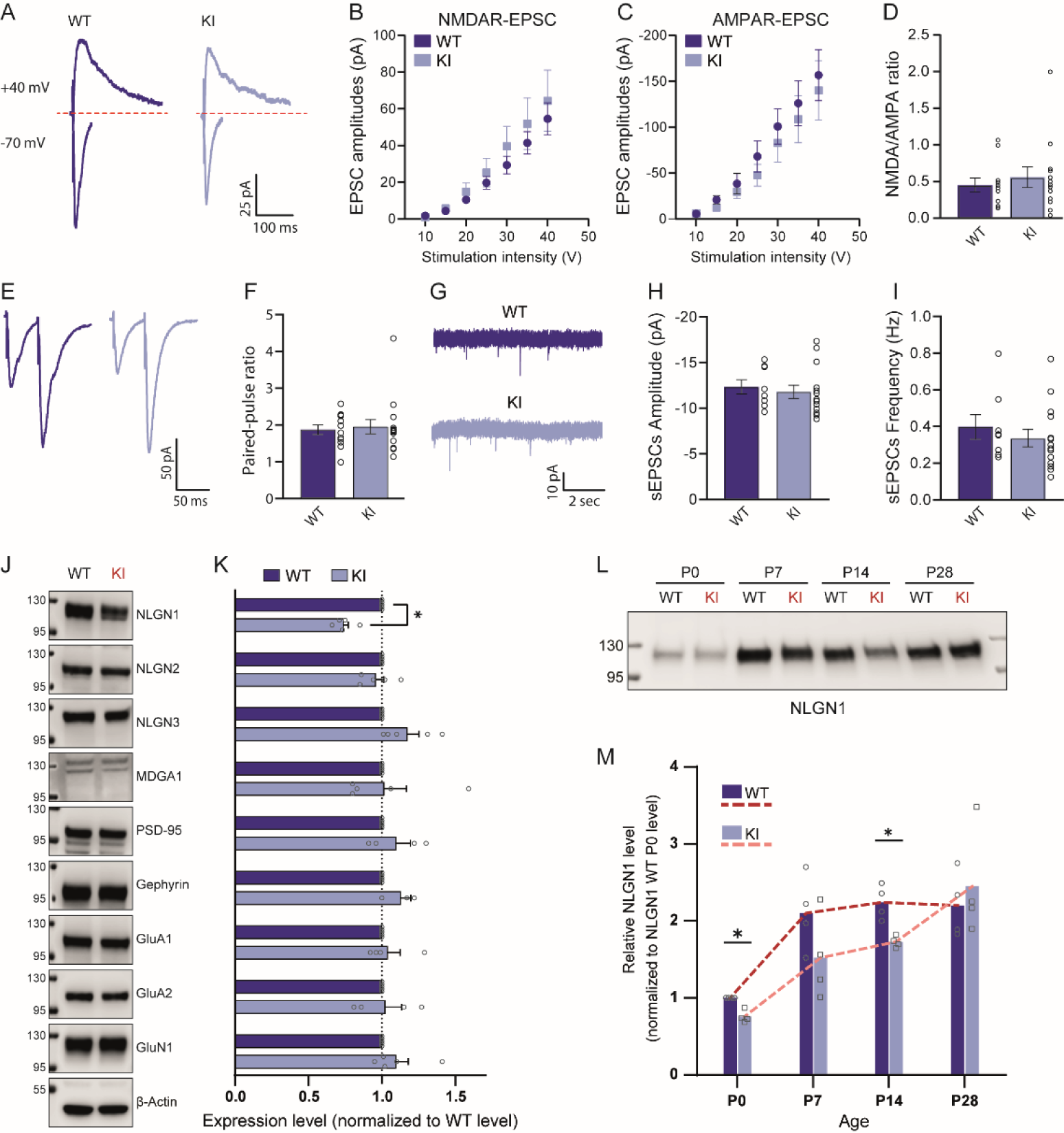
Normal synaptic transmission but decreased NLGN1 expression in KI mice. **(A)** Representative voltage-clamp recordings performed on CA1 neurons in acute slices from WT or KI mice, upon stimulation of Schaffer’s collaterals. EPSCs were recorded sequentially in the same neuron at two different holding potentials (-70 mV and +40 mV) to isolate AMPA- and NMDA-receptor mediated synaptic transmission, respectively (the NMDA-receptor dependent response is calculated at 100 ms, after the AMPA-receptor dependent response has desensitized. **(B, C)** Relationship between AMPA-receptor and NMDA-receptor mediated EPSCs and intensity of the electrical stimulation. **(D)** Paired ratio between NMDA- and AMPA-receptor mediated EPSCs measured at a 40 V stimulation. Data are presented as mean ± SEM (n = 10-12 neurons from 3 independent slices) and analyzed by unpaired t-tests (no significant difference). **(E)** Examples of AMPA-receptor mediated EPSCs in response to two stimulations separated by 50 ms in slices from WT and KI mice, respectively. **(F)** Corresponding paired pulse Ratio. **(G)** Representative electrophysiological recordings of AMPA-receptor mediated spontaneous EPSCs in CA1 neurons from WT and KI mice, respectively. **(H, I)** Bar graphs showing respectively the amplitude and frequency of those events in the two genotypes. **(J)** Representative immunoblots of hippocampal cell lysates from WT and KI mice. β-actin was used as loading control. **(K)** Graph showing the relative levels of various synaptic proteins, analyzed by semi-quantitative immunoblotting (n = 4 to 5 independent cultures for all experimental groups). Data represent mean ± SEM and were compared by 2-way ANOVA test followed by Dunnett multiple comparison test (*P < 0.05). **(L)** Western blot of proteins extracted from cortical tissue of WT or KI mice at different developmental ages (P0 to P28), separated by SDS-PAGE, and probed with NLGN1 antibody (molecular weight markers in kDa indicated on the left). Total protein staining (not shown) was used as a loading control. **(M)** Corresponding NLGN1 expression level at each developmental time. Dots in the bars represent individual animals (n = 4), from 2 independent experiments. Data represent the mean value and were compared by a multiple Mann-Whitney test (*P < 0.05). For each genotype, the evolution of NLGN1 expression over developmental time is indicated as a dotted line.

### Biochemical analysis of synaptic protein expression in KI mice

To assess the levels of various post-synaptic proteins classically associated with NLGN1, we processed lysates of hippocampal cultures from WT and KI mice at DIV 14 by SDS-PAGE, followed by immunoblotting with antibodies to NLGN1, NLGN2, NLGN3, PSD-95, gephyrin, MDGA1, the NMDA receptor subunit GluN1, and the AMPA receptor subunits GluA1 and GluA2 **(Fig. 3J)**. When normalized to actin bands on the same blots, all post-synaptic proteins showed comparable levels in cultures from WT and KI mice, except for bAP-NLGN1 which exhibited a 26 ± 3 % decrease in protein level compared to native NLGN1 from WT mice **(Fig. 3K)**. Importantly, no vertical shift appeared in the band representing bAP-NLGN1 as compared to endogenous NLGN1 in cultures from WT mice, as expected from the minor molecular weight addition brought by the bAP tag and linker (20 aa = 2 kDa), indicating a preserved integrity of the protein. To assess whether the decrease in global bAP-NLGN1 level was also present in the membrane fraction, we performed surface biotinylation of membrane proteins, followed by streptavidin pull-down and immunoblotting of NLGN1. When normalized to total protein amount, the surface level of NLGN1 was also decreased by 29 ± 2% in KI compared to WT cultures **(Fig. S5)**. Finally, to check whether the lower bAP-NLGN1 level was linked to the culture preparation, we collected fresh cortical tissue from WT and KI mice at various ages (P0, P7, P14, and P28 after birth), solubilized proteins, and performed Western blots against NLGN1 **(Fig. 3L)**. The NLGN1 level was again 25 ± 4% lower in KI mice compared to WT mice at P0 (corresponding to the difference seen in hippocampal cultures), but gradually reached the same level as in WT mice as animals grew older **(Fig. 3M)**.

Having shown in this first part of the study that the replacement of native NLGN1 by bAP- NLGN1 does not alter significantly mouse behavior, brain and synapse development, synaptic transmission, and the expression levels of common synaptic markers except for a modest reduction in NLGN1 levels, we went on to use a targeted biotinylation strategy combined with high-affinity streptavidin conjugates to selectively isolate and visualize bAP-NLGN1, as a proxy to endogenous NLGN1.

### Specific pull-down of bAP-NLGN1 and protein partners

To biotinylate bAP-NLGN1, we infected primary hippocampal neurons from KI mice at DIV 3-5 with adeno-associated viruses (AAVs) coding for the ER resident enzyme biotin ligase (BirA^ER^-IRES-GFP, called BirA+) (43), which covalently attaches exogenously supplied biotin to a central lysine residue in the bAP tag (42) **(Fig. 4A)**. As controls, we used cultures from KI mice infected with an AAV coding for GFP alone (IRES-GFP, called BirA-), and neurons from WT mice infected with BirA+ **(Fig. 4B)**. We lysed the cells at DIV 14, isolated biotinylated proteins from the lysates using streptavidin beads, and performed SDS-PAGE and immunoblots to GFP, HA, or NLGN1. GFP was detected in the starting material of all samples, while the HA tag on BirA^ER^ was detected only in the BirA+ condition, as expected **(Fig. 4C)**. NLGN1 was pulled down very efficiently by streptavidin beads from bAP-NLGN1 neurons infected by BirA+, but not in the three other conditions, showing a highly selective isolation of biotinylated NLGN1 **(Fig. 4D)**. Indeed, the NLGN1 level present in the flow through was only 22% of its initial level in the starting material **(Fig. S6)**. We then used this property to pull down proteins associated to NLGN1, including previously reported extracellular and intracellular binding partners. We detected large amounts of endogenous NLGN3 associated to bAP-NLGN1 **(Fig. 4E)**, most likely revealing the formation of NLGN1-NLGN3 heterodimers, as already reported (3, 45). bAP-NLGN1 also pulled significant amounts of PSD-95 **(Fig. 4E)**, as expected from its well characterized C-terminal interaction with PSD-95 at both biochemical and immunocytochemical levels (11, 13, 34). More surprisingly, gephyrin was also consistently found in bAP-NLGN1 pull-downs **(Fig. 4E)**, confirming that gephyrin can interact with endogenous NLGN1 in neurons as initially predicted from in vitro assays (15, 16).

**Figure 4.**
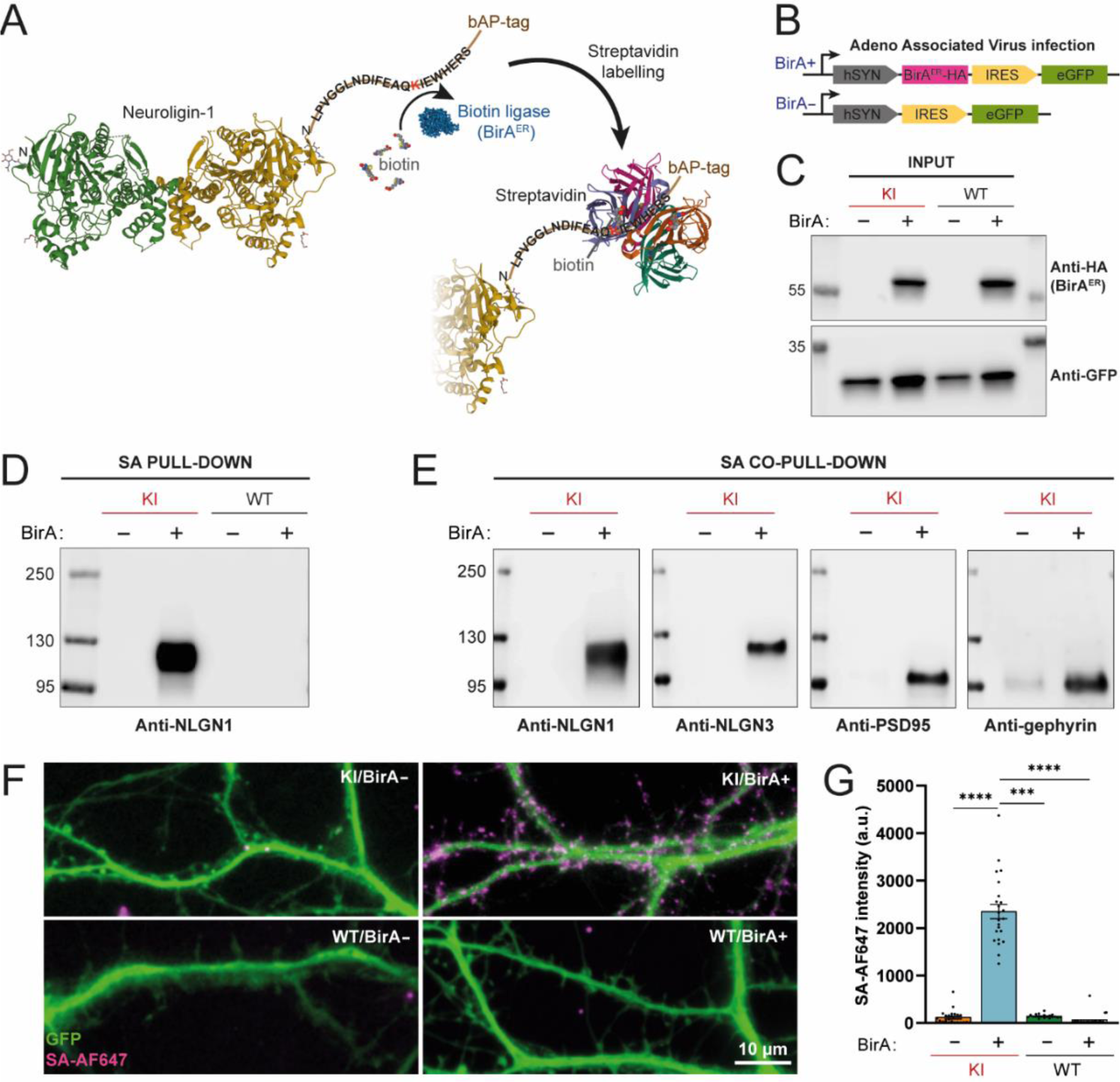
Selective isolation and detection of biotinylated bAP-NLGN1 in hippocampal cultures. **(A)** Schematic representation of NLGN1 containing the 15 aa N-terminal bAP-tag that can be enzymatically biotinylated by the ER-resident biotin ligase (BirA^ER^), and subsequently labeled with streptavidin conjugates. **(B)** Cultures from WT or KI mice were infected at DIV 3-5 with the BirA^ER^-HA-IRES-GFP virus (BirA+), or the IRES-GFP control virus (BirA-). **(C-E)** Neuron cultures were lysed at DIV 14, and protein extracts were either loaded as starting material or precipitated with streptavidin beads. Separated proteins (HA, GFP, NLGN1, NLGN3, PSD-95, gephyrin) were identified by Western blot with respective antibodies (molecular weight markers in kDa indicated on the left). **(F)** Neuron cultures were live-labeled with AF647-conjugated streptavidin (SA-AF647) and visualized by epifluorescence microscopy. Representative merged images of dendritic segments showing the GFP reporter from BirA+ or BirA- infection (green), and NLGN1 labeling with SA-AF647 (magenta). **(G)** Quantification of the SA-AF647 fluorescence intensity in each culture condition. Dots in the bars represent individual cells, with n > 20 cells for each condition, from 3 independent experiments. Data represent mean ± SEM and were compared by a Kruskal–Wallis test followed by Dunn’s multiple comparison test (***P < 0.001, ****P < 0.0001).

### bAP-NLGN1 is localized at both excitatory and inhibitory synapses

The efficiency and selectivity of the biotinylation reaction allowed us to examine in detail the localization of bAP-NLGN1 at the subcellular level, using both conventional and super-resolution microscopy. To this aim, we infected dissociated hippocampal neurons from WT or KI mice at DIV 3-5 with BirA+ or BirA- AAVs, and live stained the cultures with Alexa-647-conjugated streptavidin (SA- AF647) **(Fig. 4F,G)**. Strikingly, only neurons from KI mice infected with BirA+ showed significant SA- AF647 staining, while neurons infected with BirA- showed minimal staining, as low as neurons from WT mice infected with BirA+. Similar levels of SA-AF647 staining were observed in neurons electroporated with the BirA^ER^ plasmid and in neurons infected by BirA+ AAVs, suggesting that BirA^ER^ maximally biotinylates bAP-NLGN1. Moreover, electroporation of neurons from KI mice with a previously reported shRNA to NLGN1 (34) resulted in a 63 ± 3% decrease in SA-AF647 staining compared to the expression of a control shRNA **(Fig. S7)**, demonstrating that streptavidin specifically stains endogenous bAP-NLGN1. Streptavidin-labeled bAP-NLGN1 was enriched in puncta localized in dendritic spines, most-likely corresponding to excitatory post-synaptic densities, but also found in more elongated structures throughout the dendritic shaft, seemingly corresponding to inhibitory post- synapses. To confirm this dual localization of bAP-NLGN1, we co-electroporated neurons from KI mice with BirA^ER^ and intrabodies to either PSD-95 (46) or gephyrin (47), fused respectively to mRuby2 and GFP **(Fig. 5A)**, and quantified the fraction of bAP-NLGN1 colocalizing with those markers in the same cells **(Fig. 5B-D)**. In parallel, we checked by counter immunostaining that these intrabodies specifically labelled endogenous PSD-95 and gephyrin, respectively **(Fig. S8)**. Strikingly, bAP-NLGN1 accumulated as strongly at PSD-95 and gephyrin puncta (enrichment in post-synapse versus shaft around 3), and the proportion of PSD-95 or gephyrin puncta that contained bAP-NLGN1 were respectively 70 ± 2% and 54 ± 3% **(Fig. 5E,F)**, showing that more than half inhibitory post-synapses actually recruit NLGN1. The proportion of bAP-NLGN1 within PSD-95 and gephyrin puncta was 57% and 15%, respectively **(Fig. 5D)**, in relation to the fact that there are around 2 times more excitatory synapses than inhibitory synapses in those cultures **(Figs. S4A-C and S8C)**. There was also a 22 % fraction of extra-synaptic bAP- NLGN1 clusters **(Fig. 5C,D),** significantly smaller than synaptic bAP-NLGN1 clusters **(Fig. 5G)**, and a 6 % fraction of bAP-NLGN1 that co-localized with both PSD-95 and gephyrin **(Fig. 5C,D)**, likely corresponding to doubly innervated dendritic spines (48).

**Figure 5.**
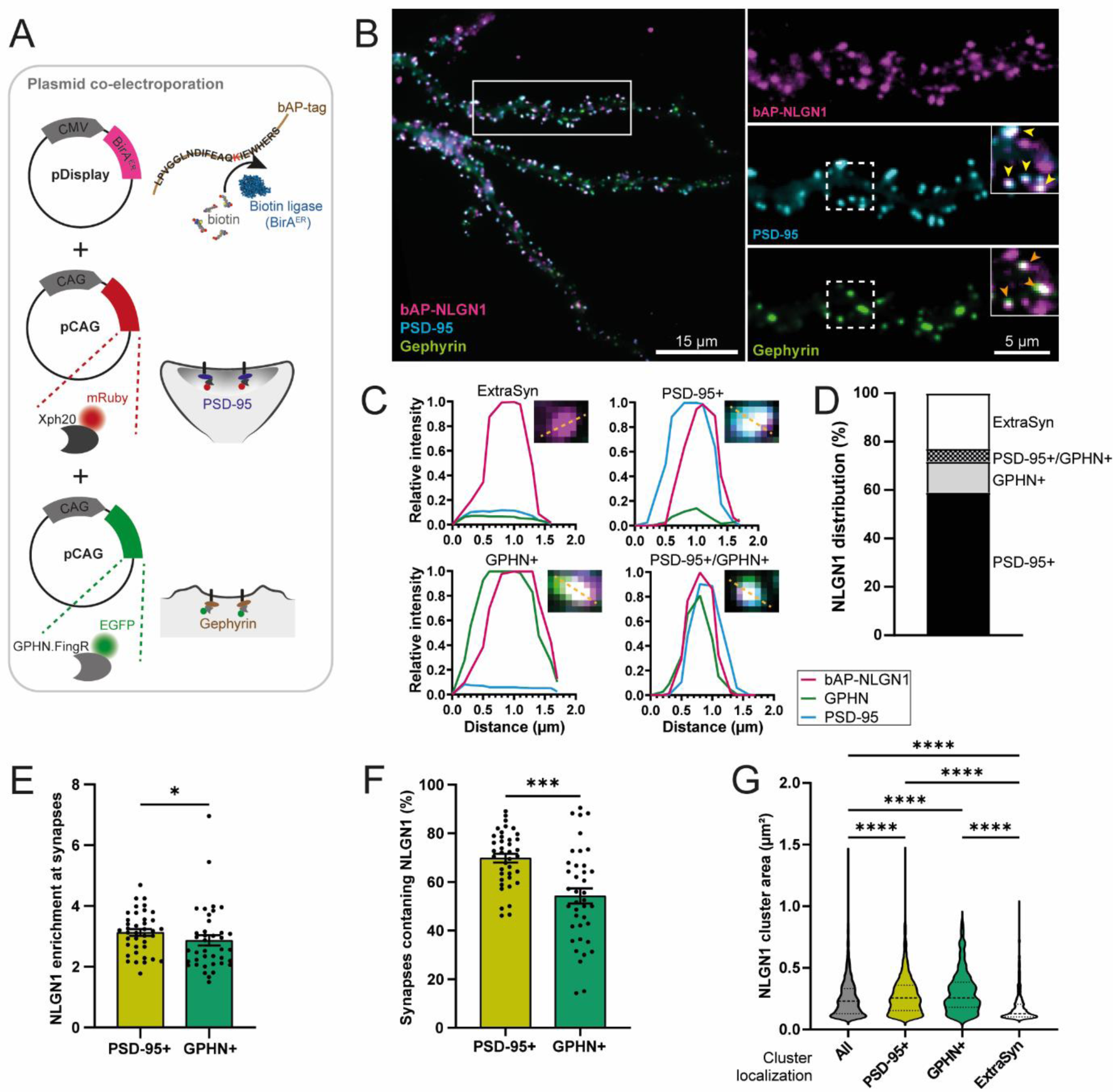
Sub-cellular localization of biotinylated bAP-NLGN1 at excitatory and inhibitory synapses. **(A)** Dissociated neurons from bAP-NLGN1 hippocampal cultures were co-electroporated at DIV 0 with plasmids encoding BirA^ER^ and two intrabodies (Xph20-mRuby2 and GPHN.FingR-GFP), which label respectively the excitatory and inhibitory post-synaptic proteins PSD-95 and gephyrin. The BirA^ER^ plasmid has no reporter. **(B)** At DIV 14, neurons were live-labeled with SA-AF647 and visualized by epifluorescence microscopy. Representative image of a neuron showing bAP-NLGN1 labeled with SA-AF647 (magenta), PSD-95 positive synapses (cyan), and gephyrin positive synapses (green). An example of dendritic segment corresponding to the rectangle area is shown on the right. Insets showing zoomed synapses, with examples of SA-AF647 clusters overlapped with PSD- 95 (yellow arrows), or gephyrin (orange arrows) positive puncta. **(C)** Representative line scans across synaptic ROIs (insets) showing NLGN1 clusters either in an extrasynaptic location (ExtraSyn), overlapping with PSD-95 (PSD-95+), overlapping with gephyrin (GPHN+), or with both PSD-95 and gephyrin (PSD-95+/GPHN+). **(D)** Representation of bAP-NLGN1 cluster distribution throughout the dendrite. **(E, F)** Bar plots representing the enrichment of bAP-NLGN1 at PSD-95 or gephyrin positive puncta, and the fraction of these puncta overlapping with bAP-NLGN1, respectively. Dots in the bars represent individual cells, with n = 39 cells for each condition, from 2 independent experiments. Data represent mean ± SEM and were compared by a Mann-Whitney test (*P < 0.05, ***P < 0.001). **(G)** Violin plots showing the surface area of individual bAP-NLGN1 clusters as a function of their localization. Quantification was performed either on all NLGN1 clusters present throughout the dendritic segment (All), on clusters overlapping with PSD-95 or gephyrin (PSD-95+/GPHN+), or on clusters localized at extrasynaptic sites (ExtraSyn) (n = 436-2224 clusters in 39 cells, from 2 independent experiments). Data were compared by a Kruskal–Wallis test followed by Dunn’s multiple comparison test (****P < 0.0001).

### bAP-NLGN1 forms nanodomains in both excitatory and inhibitory post-synapses

We further examined the nanoscale localization of biotinylated bAP-NLGN1 stained with SA-AF647 using direct STochastic Optical Reconstruction Microscopy (dSTORM) (10, 27, 28), in relation to excitatory or inhibitory post-synapses labeled with the specific intrabodies described above **(Fig. 6A,B)**. Super-resolved maps integrating all single molecule localizations revealed a diffuse distribution of bAP-NLGN1 in the dendritic shaft, as well as a specific accumulation in excitatory and inhibitory post-synapses, with a similar synaptic versus shaft enrichment around 4 **(Fig. 6C)**. We then analyzed the sub-synaptic distribution of bAP-NLGN1 using a segmentation approach based on Voronoï polygons (49) **(Fig. 6D,E)**. bAP-NLGN1 was found to form several small domains (between 0 and 5, average around 1) in both excitatory and inhibitory post-synapses **(Fig. 6F)**, and whose size ranged between 30-100 nm **(Fig. 6G)**. Those nanodomains were highly enriched, i.e. reaching a density of single molecule detections 80-fold larger than that in the shaft **(Fig. 6H,J)**. Strikingly, the number of bAP-NLGN1 nanodomains **(Fig. 5I,K)**, but not their size **(Fig. S9C,D)**, positively correlated with the projected area of the corresponding PSD or gephyrin puncta. By normalizing the integrated number of single molecule localizations in post-synapses by the corresponding number of single SA-AF647 molecules isolated on a coverslip and imaged under the same conditions **(Fig. S9E-J)**, we computed that 10.0 ± 0.6 SA-AF647 molecules were bound to biotinylated bAP-NLGN1 in excitatory post- synapses (mean ± SEM from 1139 PSD-95 positive puncta), and 11.3 ± 0.9 SA-AF647 molecules in inhibitory post-synapses (from 631 gephyrin positive puncta). The numbers of SA-AF647 within bAP- NLGN1 nanodomains located inside PSD-95 or gephyrin puncta were 5.3 ± 0.3 and 4.7 ± 0.3, respectively. Increasing the concentration of SA-AF647 did not improve the specific fluorescence signal on neurons expressing BirA^ER^, suggesting that biotinylated bAP-NLGN1 binding sites are saturated by streptavidin.

**Figure 6.**
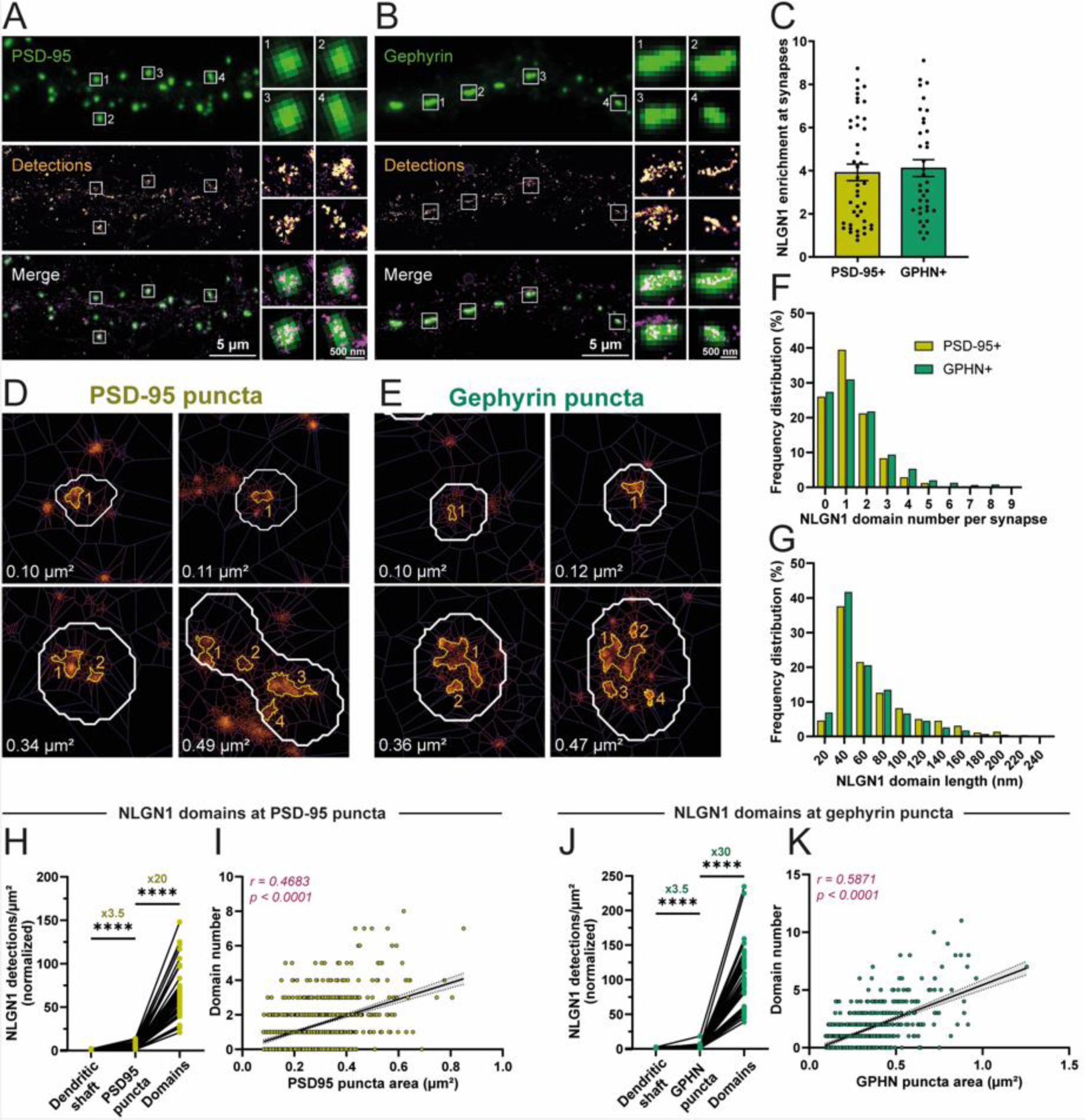
Super-resolved localization of biotinylated bAP-NLGN1. Dissociated neurons from bAP-NLGN1 hippocampal cultures were electroporated at DIV 0 with a combination of BirA^ER^ and either Xph20-GFP or GPHN.FingR-GFP. dSTORM experiments were performed at DIV 14 on fixed cultures, after live-labeling neurons with SA-AF647. **(A, B)** Representative images of dendritic segments showing PSD-95 or gephyrin positive synapses (green), the super-resolved localization map of all bAP-NLGN1 detections (gold), and merged images (synaptic puncta in green and detections in magenta). On the right, individual examples of excitatory and inhibitory synapses containing bAP-NLGN1 localizations. **(C)** Bar plots representing the enrichment of bAP-NLGN1 localizations at PSD-95 or gephyrin positive puncta. Dots in the bars represent individual cells. Data represent mean ± SEM and were compared by a Mann-Whitney test (*p* = 0.53). **(D, E)** Representative images of bAP-NLGN1 localization obtained with SR-Tesseler, showing the presence of a variable number of nanodomains (outlines in gold) within PSD-95 and gephyrin positive puncta, respectively (outlines in white). Note that the number of nanodomains increases with the surface area of the post-synaptic puncta (indicated in white). **(F, G)** Frequency distribution of the number and length, respectively, of bAP-NLGN1 nanodomains within PSD-95 and gephyrin positive puncta. **(H, J)** Average surface density of bAP-NLGN1 detections throughout the dendritic shaft, in post-synapses, or in synaptic nanodomains, for either PSD-95 or gephyrin positive puncta. Dots represent individual cell at each location. Data were compared by a Kruskal– Wallis test followed by Dunn’s multiple comparison test (****P < 0.0001). **(I, K)** Correlation between the bAP- NLGN1 domain number, and the PSD-95 or gephyrin post-synapse surface area, respectively. Dots represent individual synaptic puncta. r, Pearson’s correlation coefficient; black line, linear regression with 95% confidence interval (grey). All data related to PSD-95+ puncta were obtained from 4 independent experiments, with 1390 domains / 1065 synaptic puncta / 43 cells analyzed. Data related to GPHN+ puncta were obtained from 3 independent experiments, with 984 domains / 631 synaptic puncta / 36 cells analyzed.

### Nanogold labeling of bAP-NLGN1 also reveals nanodomains in the synaptic cleft

Finally, we sought to visualize the distribution of endogenous bAP-NLGN1 in the synaptic cleft at an even higher spatial resolution using TEM, which also has the advantage of highlighting membrane ultrastructure. Since 2D dissociated cultures are flat and form relatively dispersed synapses, they are not an ideal sample for TEM. Thus, we turned to organotypic hippocampal slices, a preparation with well-preserved neuronal architecture and synaptic connectivity (22, 50). Slices from either WT or KI mice were infected with BirA^+^ at DIV 3 and used at DIV 14 **(Fig. 7A)**. A first set of organotypic slices was incubated live with SA-AF647, then fixed and observed in confocal microscopy **(Fig. 7B-D)**. There was a specific SA-AF647 signal on slices from KI - but not WT mice-, preferentially localized in the CA3 region which might reflect a better sensitivity to the AAV infection **(Fig. S10)**. Moreover, in slices from KI mice, there was a specific accumulation of bAP-NLGN1 in the different synapse types formed on proximal and distal dendrites of CA3 pyramidal neurons **(Fig. 7E,F)**. A second set of organotypic slices was incubated live with streptavidin conjugated to a 1.4 nm gold nanoparticle, then fixed, embedded in resin, and cut in ultra-thin sections before observation in TEM. In cultures from KI mice infected with BirA^+^, there was a striking accumulation of multiple silver particles that decorated the synaptic cleft **(Fig. 7H,J)**, while this type of signal was absent in WT slices also treated with BirA+ **(Fig. 7G,I)**. Some non-specific silver aggregates were also visible throughout the tissue, but they were not adjacent to synapses. Approximatively 15% of synapses detected by TEM contained silver particles, which may be due to a combination of factors: i) not all neurons are infected by BirA^+^; ii) not all synapses contain bAP-NLGN1; and iii) the thin 70-nm sections cut for TEM may miss some silver particles that are actually present in a 300-nm wide PSD (51). The number of silver particles in the synaptic cleft was on average of 8.2 ± 1.0 **(Fig. 7K)**, close to the values found in dSTORM **(Fig. S9)**. Finally, silver particles were not distributed homogeneously in the synaptic cleft, but formed several sub-domains that strikingly resembled those seen in dSTORM **(Fig S6)**. The number of nanodomains was again between 1 to 4 **(Fig. 7L)**.

**Figure 7.**
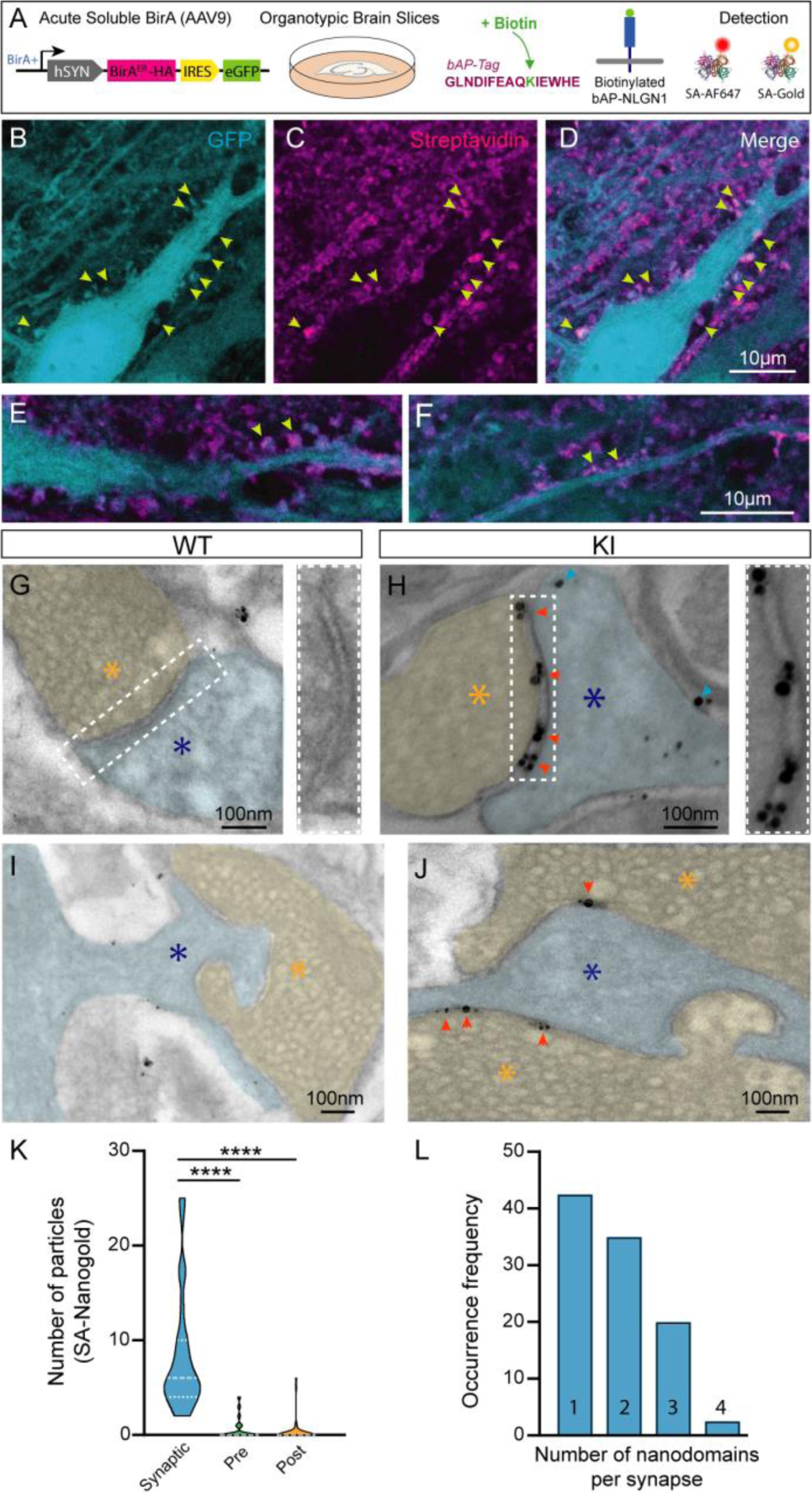
Visualization of bAP-NLGN1 in the synaptic cleft by TEM. **(A)** Molecular strategy to biotinylate bAP-NLGN1 in organotypic slices and stain it with fluorescent or nanogold streptavidin conjugates. **(B**-**D)** Representative images showing a CA3 neuron from a KI organotypic slice infected with hSYN-BirA^ER^-IRES-eGFP (cyan), where bAP-NLGN1 was labeled with SA-AF647 (magenta). **(E**, **F)** Higher magnification images of synapses located on proximal and distal dendrites, respectively (yellow arrows). **(G-J)** Representative TEM images of synapses in the CA3 region of organotypic slices from WT and KI mice, respectively. Slices were infected with hSYN-BirA^ER^-IRES-eGFP at DIV 3, left in the incubator for 2 weeks, then incubated with SA-Nanogold, fixed, and silver enhanced. Pre- and post-synapses are highlighted in yellow and blue, respectively (with stars of the corresponding color). Images in **(G**, **H)** correspond to canonical synapses formed by CA3 axons onto distal dendrites, while images in **(I**, **J)** likely correspond to giant synapses formed between mossy fibers and thorny excrescences on proximal dendrites. Insets on the right show zoom on the synaptic cleft in the two genotypes. Red and blue arrows show the location of synaptic and extrasynaptic endogenous NLGN1, respectively. **(K)** Subcellular distribution of silver enhanced SA-nanogold particles in the synaptic cleft and in pre-synaptic or post-synaptic compartments. Data represent mean ± SEM and were compared by a Brown-Forsythe test followed by Dunn’s multiple comparison test (****P < 0.0001). **(L)** Occurrence frequency of the number of bAP-NLGN1 nanodomains labeled with SA-Nanogold in the synaptic cleft (n = 40 synapses from 3 different experiments were analyzed).

## Discussion

Advances in our understanding of synaptic connectivity in the brain require a combination of novel animal models and experimental approaches, giving access to the targeting of endogenous neuronal adhesion proteins with high selectivity and spatial resolution. In this study, we have generated and extensively characterized a new transgenic mouse strain in which endogenous NLGN1 was tagged with a small peptide (bAP) allowing for its functional biotinylation by exogenously delivered biotin ligase, followed by highly efficient isolation and visualization with various streptavidin conjugates. With this tool in hand, we reveal that native NLGN1 can bind both PSD-95 and gephyrin at the biochemical level, and distributes equally well at excitatory and inhibitory synapses at the fluorescence microscopy level. Using super-resolution light microscopy and electron microscopy, we further highlight that bAP- NLGN1 is organized in small nanodomains in the synaptic cleft, whose number scales with the post- synapse size.

By considering a variety of behavioral parameters (body weight, size, fertility, locomotion, anxiety, …), and judging from the gross anatomy and cell density in different brain structures (cortex, hippocampus), KI mice were comparable to control WT mice of the same genetic background. The density of excitatory and inhibitory synapses quantified in either acute hippocampal slices by immunofluorescence staining of VGlut1 and VGAT, or in hippocampal cultures by labeling of gephyrin and PSD-95, was similar in WT and KI mice. The synaptic ultrastructure evaluated by TEM as well as the AMPA- and NMDA-receptor dependent synaptic transmission measured by electrophysiology were also similar in WT and KI mice. The levels of most post-synaptic proteins isolated from hippocampal cultures and revealed by Western blot were comparable between WT and KI genotypes, with the exception of bAP-NLGN1 itself whose level was reduced by 26 ± 3 % with respect to NLGN1 from WT mice, in both total and surface protein samples. For comparison, 10%, 50%, and 60% reductions in the protein levels of HA-NLGN1, bAP-GluA2, and SEP-GluA1 were observed in brain lysates from their respective knock-in mice (39, 43, 52), suggesting that the introduction of an external tag in a gene of interest tends to reduce the protein level in the KI mouse. Interestingly, the larger the tag, e.g. GFP, bAP, HA (238, 15, and 9 aa, respectively), the stronger the reduction in protein level. Specifically, although the bAP tag is the same in our bAP-NLGN1 KI mouse model and in a recently published bAP-GluA2 KI mouse (43), the addition of 48 amino acids encoding the bAP-TEV and linker sequence on GluA2 (5 kDa) was larger than the overall sequence we used here for bAP-NLGN1 (21 amino acids, 3 kDa), such that the band corresponding to bAP-NLGN1 was barely above that of endogenous NLGN1. The reduction in bAP-NLGN1 protein level can be explained by the fact that the introduction of the bAP-tag in the NLGN1 gene perturbs the transcription machinery, resulting in reduced mRNA level, as observed in the SEP-GluA1 KI (52), or by the reduced stability of the mRNA transcript, which translates into reduced protein translation. Nevertheless, the amount of bAP-NLGN1 measured in tissue from P0-P30 old KI mice progressively reached the levels in WT mice, suggesting a developmental adjustment. In any case, the number and ultrastructure of both excitatory and inhibitory synapses in acute slices or cultures from KI mice were identical to those from WT controls, in line with the observation that synapses are generally unaffected in constitutive NLGN1 KO mice (44). There was no significant change in NLGN2 or NLGN3 protein levels in the bAP-NLGN1 KI mouse, indicating no compensation by other NLGN isoforms, in contrast to the constitutive NLGN1 KO which displays increased NLGN3 levels (53) or to the SEP-GluA1 KI which exhibits increases in GluA2/GluA3 expression (52).

Based on organotypic and primary hippocampal cultures from KI mice combined with cell-specific biotinylation provided by the expression of BirA^ER^, we were able to isolate and label bAP-NLGN1 with high selectivity using streptavidin conjugates. Indeed, no NLGN1 was pulled-down by streptavidin beads or fluorescently labeled by SA-AF647 in cultures from WT mice expressing BirA^ER^, or in cultures from KI mice not expressing BirA^ER^. Moreover, knocking down endogenous NLGN1 with a previously published shRNA (34) decreased the surface SA-AF647 signal on neurons from KI mice co-expressing BirA^ER^ as compared to a control shRNA, demonstrating that streptavidin specifically labels native bAP- NLGN1 and further validating this shRNA to NLGN1. We previously managed to live label endogenous NLGN1 with purified β-NRXN1-Fc (13, 28), but the staining was quite weak compared to the streptavidin labeling obtained here, most likely due to an enormous difference in binding affinity. Streptavidin pull-down of biotinylated bAP-NLGN1 was highly efficient as it captured roughly 80% of NLGN1 proteins from cell lysates, a level that we never reached with antibodies to NLGN1 (10, 16). Interestingly, whereas β-NRXN1-Fc and MDGA1/2-Fc were used to pull-down protein binding partners (7, 54), to our knowledge no one ever reported the use of NLGN1-Fc as a bait, potentially because of the poor stability of this purified protein (55). The streptavidin pull-down of bAP-NLGN1 allowed the identification of already reported binding partners, including PSD-95 which can interact with the C-terminal PDZ domain binding motif of NLGN1 (11) and NLGN3 which is likely to form heterodimers with NLGN1 through an extracellular coiled coil interface (3, 4, 56). Interestingly, although gephyrin is well-known to interact with NLGN2 (15, 32), it was pulled down efficiently by bAP-NLGN1. Yeast two-hybrid and peptide pull-down assays had previously shown significant binding between the gephyrin E domain and NLGN1 in vitro (15, 16), but this interaction had never been demonstrated in neurons. The coupling between endogenous bAP-NLGN1 and gephyrin likely involves the highly conserved motif (PPDYTLAMRRSPDDIP) located in the middle of the NLGN1 intracellular domain, whose binding to gephyrin is inhibited by tyrosine phosphorylation (16, 22). Based on the expression of NLGN1 tyrosine point mutants (Y782A/F) that mimic phosphorylated or non-phosphorylated states, we originally proposed that NLGN1 could be localized at either excitatory or inhibitory synapses, depending on the NLGN1 tyrosine phosphorylation level (16, 22), but the actual localization might have been biased by NLGN1 over-expression. Here, we clearly demonstrate that native bAP-NLGN1 is similarly enriched at both excitatory and inhibitory synapses, challenging the historical view that NLGN1 is exclusively localized at excitatory synapses (31). Apart from the phosphorylation mechanism, it is possible that different splice variants of NLGN1 are present at excitatory and inhibitory synapses where they would interact with specific NRXNs, e.g. NLGN1 with site B being preferentially targeted to excitatory synapses versus NLGN1 w/o site B at inhibitory synapses, although there are modest amounts of mRNA coding for NLGN1(B-) in rodent brain tissue (36–38). Interestingly, a recent study that improved the immunoreactivity of antibodies to several synaptic proteins using an alternative fixative to paraformaldehyde (glyoxal), showed equivalent immunofluorescence signals of endogenous NLGN1 at both excitatory and inhibitory synapses in brain slices (57).

The cell-specific biotinylation procedure allowed us to image bAP-NLGN1 at the cell surface by super-resolution microscopy (dSTORM) after live staining with SA-AF647. First, there was a significant reservoir of non-synaptic bAP-NLGN1 molecules, which might correspond to the diffusive bAP-NLGN1 molecules previously identified by single molecule tracking on moderately over-expressed bAP-NLGN1 (10, 28). Second, endogenous bAP-NLGN1 molecules were enriched approximately 3-fold in both PSD- 95 and gephyrin puncta as compared to the dendrite shaft. Third, within PSD-95 rich puncta, bAP- NLGN1 formed a small number of nanodomains of average size 50 nm and containing very high protein density. The number (but not the area) of such nanodomains was positively correlated with the surface area of the post-synapse, as reported in brain slices from HA-NLGN1 KI mice immunostained with anti-HA antibodies and processed for expansion microscopy (40). Given that streptavidin is tetravalent and was used at a concentration that saturates biotin binding sites on the neuron surface, one SA-AF647 molecule likely binds two bAP-NLGN1 dimers. Thus, based on the number of streptavidin molecules counted in dSTORM and TEM, we give an estimate of 16-20 bAP-NLGN1 dimers per synapse. The actual number of synaptic NLGN1 dimers is likely to be in the range of 22-27, since surface bAP-NLGN1 levels are 26% lower than those of endogenous NLGN1, which is close to the estimate of 21 copies of NLGN3 molecules made by quantitative mass spectrometry (58). Interestingly, bAP-NLGN1 also formed nanodomains at inhibitory post-synapses, with a similar correlation between the number of domains and the area of gephyrin puncta. Although we did not resolve them here, the scaffolding molecules PSD-95 and gephyrin themselves are known to form substructures with high protein density in the post-synapse, but tend to form larger clusters than bAP-NLGN1 nanodomains (59–62). This observation suggests that bAP-NLGN1 obeys an internal organization which is dictated not only by interactions with the post-synaptic scaffold, but also potentially via trans-synaptic interactions with pre-synaptic NRXNs, which also forms nanodomains of similar size (28, 63, 64). Recently, NLGN1 was shown to partition in high-order liquid condensates together with PSD-95 and the AMPA receptor auxiliary subunit stargazin (65, 66). This process might also underlie the organization of bAP-NLGN1 in nanodomains seen both at the dSTORM and TEM levels. Similarly, it was shown that gephyrin forms condensates with the cytoplasmic loops of glycine or GABA_A_ receptors (66). Given the similarity of organization of bAP-NLGN1 at both excitatory and inhibitory synapses, an interesting perspective would be to evaluate whether gephyrin also phase separates with binding motifs of the NLGN1 intracellular domain (16, 67).

As a conclusion, this tagged transgenic mouse strain offers the possibility to biochemically isolate endogenous NLGN1 with high yield and thus potentially identify new binding partners, and to image NLGN1 with high selectivity in single neurons at the super-resolution level. Given that it has been notoriously difficult to generate efficient antibodies to many neuronal cell adhesion molecules including NRXNs, NLGNs, and LRRTMs, it might be appealing in the future to combine several knock-in approaches with orthogonal labeling strategies to visualize those trans-synaptic complexes in multi- color imaging.

## Materials and Methods

### Generation and amplification of the bAP-NLGN1 transgenic mouse line

A new transgenic mouse strain was generated in which the endogenous *Nlgn1* gene in chromosome 3 was N-terminally flanked by *<u>a small linker</u>* and a DNA sequence coding for the 15 aa **biotin acceptor peptide** (bAP) (42), *<u>LPV</u>***GLNDIFEAQKIEWHE***<u>RS</u>*, inserted after the signal peptide. The generation of this mouse strain was made at the PHENOMIN-ICS (Strasbourg, France; http://www.phenomin.fr) and was based on the CRISPR/Cas9 strategy targeting exon1 of the *Nlgn1* gene (**Fig. S1**). Using the CRISPOR software (http://crispor.tefor.net/), a specific guide RNA of 19 nucleotides (gRNA79, 5’- TCACGTACTCTCTCAAAAGT -3’) was designed so as to produce a double strand break of the *Nlgn1* gene at a PAM motif (TGG) ideally located at 2 bp from the end of the signal peptide sequence. The bioinformatics analysis gave the following parameters: specificity score 79; on-target efficiency 44; off-target efficiency 37; out-of-frame score 75. There was no predicted off-target gene with 0, 1, or 2 mismatches; there were 13 off-target genes with 3 mismatches and 78 with 4 mismatches, which targeted primarily introns or intergenic sequences. The guide RNA gRNA79 was produced at PHENOMIN-ICS (68), while the Cas9 protein and single stranded DNA (ssDNA) were purchased from Integrated DNA Technology (idtdna.com). CRISPR guide efficiency was tested *in vitro* using Sureguide kit (Agilent Technologies 5190-7716). Seventy-five fertilized oocytes from C57BL/6J mice were electroporated with purified 1 ng/µL Cas9 mRNA + 6 ng/µL gRNA79, and 25 ng/µL ssDNA, then embryos were transplanted into 4 C57BL/6J females. Thirty-two pups were born, of which 3 mosaic females F_0_ founders carrying the correct bAP insertion emerged. The sequence of the bAP was confirmed by Sanger sequencing. This founder was used to create a mouse colony, which was later imported in our own animal facility at the Interdisciplinary Institute for Neuroscience (IINS) and named B6J-Nlgn1^em1(bAP)Ics/Iins^. The colony was first amplified at the heterozygous level, then a homozygous strain was created. To avoid genetic drift, mice were back-crossed with C57BL/6J mice every 10 generations. Animals were regularly genotyped by polymerase chain reaction (PCR) assay on tail biopsies, with the following primers to detect *Nlgn1*: forward 5’-TGCTTTGAGCATTCCTTCACAAGCT- 3’ and reverse 5’-CCTAATCTTGCCAAAGTTAGTAGTA-3’. All animals used in this study were homozygous and WT counterparts of the same genetic background (C57BL/6J).

### Animals

All procedures involving animals and their care were conducted in accordance with the European guidelines for the car and use of laboratory animals, and the guidelines issued by the University of Bordeaux animal experimental committee (CE50; animal facilities authorizations #A33063940 and ethical project authorizations #21725-2019081114558294). Housing was at a constant temperature (21°C) and humidity (60%), under a fixed 12h light/dark cycle with food and water available ad libitum. Overall, for behavioral testing, an equal number of male and female mice were used from both WT and KI genotypes. For primary hippocampal cultures made from P0 pups, no sex determination was made, but the fact that cells isolated from several male and female animals were mixed together ensured an approximate equal repartition of sex origin. Hippocampal slices came from one male or female mouse, but the multiplication of experiments ensured randomization of sexes. Every effort was made to minimize the numbers of experimental animals.

### Behavioral testing

We used the SmithKline Beecham, Harwell, Imperial College, Royal London Hospital phenotype assessment (SHIRPA) protocol (43, 69) to assess the behavioral phenotype of bAP-NL1 mice. The SHIRPA test involves 31 parameters, which were divided into 4 groups (General aspect, Locomotion, Sensory and Anxiety). In the first stage, mice were visually examined for the skin color (white to red), member tone (no resistance to extreme resistance), abdominal tone, irritability, righting reflex, aggression and vocalization. Then, mice were initially placed inside a Perspex jar for 5 min and then evaluated for body position (from flat to repeated leaping), spontaneous activity (from immobile to extreme), spasm rate (from none to marked), and body tremor (from absent to marked). Mice were then transferred in an open-field made of clear Perspex for 10-min to record the following behavior: transfer arousal (from prolonged freezing to manic-type behavior), locomotor activity, visual placing, piloerection, gait (from normal to incapacitated), pelvic elevation (from flattened to normal), tail position (from dragging to Straub type), touch escape (from no response to vigorous attempt at escaping from being touched), fear and positional passivity. The mice were then lifted by the tail and the extent of struggling (from tail to no struggle at all) was estimated. On being lowered by the tail toward a horizontal grid, the mice were evaluated for body tone by pressing on each side (from flaccid to hard resistance), pinna (from absent to multiple flicks) and corneal (from absent to multiple blinks) reflexes. Finally, mice were gently pulled by the tail and evaluated on vertical grid for the catalepsy and negative geotaxis (turns and climb to fall down).

As the SHIRPA results suggested a possible effect on anxiety levels of KI mice, this scoring was complemented by performing the elevated plus maze test in adult WT and KI mouse line. This test consisted in a platform of four opposite arms (40 cm) two of them are open and two are closed arms (enclosed by 15 cm high walls). The apparatus was elevated at 55 cm from the floor. The task was recorded and analyzed with the software EthovisionXT (Noldus) and we measured the time spent in each arm in trials of 10 min. The luminosity of the room was 12 lux in the open arms.

### DNA constructs and viruses

The plasmid coding for Endoplasmic Reticulum (ER) resident biotin ligase, BirA^ER^ (42), was a gift from A. Ting (Stanford University, Palo Alto, CA, USA). The intrabodies to PSD-95 (70), Xph20-GFP and Xph20-mRuby2 (Addgene #135,530 pCAG_Xph20-eGFP-CCR5TC; #135,531 pCAG_Xph20-mRuby2- CCR5TC), were gifts from M. Sainlos (IINS, University of Bordeaux). The intrabody to gephyrin, GPHN.FingR-GFP (47), (Addgene # 46296 pCAG_GPHN.FingR-eGFP-CCR5TC), was a gift from D. Arnold (University of Southern California, Los Angeles, CA, USA). Short hairpin RNA to murine NLGN1, and its control shRNA to the heat shock protein p53 (34), were kind gifts from P. Scheiffele (Biozentrum, Basel, Switzerland). The plasmids coding for BirA^ER^-IRES-GFP and IRES-GFP were gifts from D. Choquet, and previously validated (43). The production of the corresponding AAVs was made by the viral core facility of the Bordeaux Neurocampus IMN. Viral titers were between 1.2 × 10^13^ and 2.0 × 10^13^ genome- containing particles (GCP)/mL.

### Immunohistochemistry on brain slices

WT and KI mice at P30 were anesthetized using Pentobarbital (300mg/Kg) and Lurocaine (30mg/Kg) injected intraperitoneally and then perfused intracardially with 50 mL of PBS followed by 50 mL of 4% PFA. The brains were extracted, placed in PFA for 48h and then in a 30% sucrose solution for 48h. After this period, brains were frozen in -30°C Isopentane for 1 min. 40 microns thick coronal sections were then cut using a cryostat (Leica CM1800) and placed in antifreeze solution at -20°C until used. For slice immunostaining, after a PBS wash, the tissue was permeabilized, nonspecific binding sites were blocked and slices were incubated overnight with a rabbit anti-VgluT1 (1:1000, Synaptic System), a mouse anti-VGAT (1:1000, Synaptic System) antibody. Primary antibodies were subsequently detected with rabbit 647-conjugated secondary antibodies (1:500, 2h incubation; Invitrogen). Slices were mounted on charged microscope slides (Superfrost/Plus, Fisher Scientific) and stored at 4°C prior to image acquisition.

### Image acquisition with confocal microscopy and analysis

All in vitro fluorescence imaging quantification analyses were performed on images acquired using a spinning-disk microscope (Leica, Germany). Images were scanned sequentially to prevent non-specific bleed-through signal using 405 and 647 nm laser excitation and a 60× (NA 1:42) oil immersion objective. For the immunohistochemical characterization of brain tissue obtained from WT and bAP- NLGN1 mice, surface and intensity for each signal were measured in a series of three different cortical and hippocampal sections ranging from bregma -1.7 to bregma –1.9 mm, with a total of 14 different spots for each hemisphere. All image quantification was performed using ImageJ (National Institute of Health [NIH]) software. A background correction was first applied at the same level for every image analyzed before any quantification. A macro developed in-house was used to perform all quantifications.

### Nissl staining on brain slices

Fixed mouse brain slices were first dehydrated into chloroform / ethanol solution (1:1) overnight at room temperature. Then, sections were progressively hydrated with gradient alcohol and stained with Toluidine blue for 20 min at 55°C and differentiated in 95% alcohol. Finally, brain sections were dehydrated with absolute ethanol for 1 min and cleared with xylene. All sections were then mounted with Permount mounting medium. Neuronal cell counting in the Cortex and the Hippocampus area CA1 was conducted using a light transmission microscope (magnification, x40, Leica, FR). For quantitative analysis, cell bodies were counted in 4 non-overlapping regions of interest (ROI) for each structure by using a homemade macro running on Image-J software.

### Ultratructural characterization of synapses in brain slices by TEM

Following i.p. injection of Pentobarbital (300mg/Kg) and Lurocaine (30mg/Kg), P30 mice were transcardially perfused with 50 ml of ice-cold sodium phosphate buffer saline (PBS; 0.1M; pH 7.4) followed by 100 mL of a mix composed of 4% paraformaldehyde and 1% glutaraldehyde diluted in PBS. Dissected brains were extracted and post-fixed overnight in 4% PFA / 1% glutaraldehyde at 4°C. Cortex from WT and KI mouse brains were manually microdissected. Sections were then washed 3 times in ddH2O and incubated on ice for 1 hour in a solution composed of 1.5% potassium ferrocyanide and 1.5% osmium tetroxide (EMS) diluted in ddH2O. Sections were dehydrated in graded ethanol and then flat-embedded in Epoxy 812 medium (Catalogue no.14121; EMS, United-States) for 48h at 60°C. Trapezoidal blocs of tissue from cortex were cut from the resin flat-embedded sections. Each piece of tissue was cut into 70 nm ultrathin sections with an ultramicrotome (EM UC7, Leica, Germany). Ultrathin sections were then collected on bare 150-mesh copper grids and examined under a transmission electron microscope (Hitachi H7650; Japan) at 80 kV. Profiles of synaptic terminals were readily identified as such by their synaptic vesicles content. Using an integrated digital camera (ORIUS SC1000 11MPx, GATAN, Ametek, United-States), axon terminals were imaged randomly, at a working magnification between 2500X to 8,000X, by acquiring an image of every such profile encountered, until 30 or more showing a full contour and distinct content were available for analysis, in each mouse.

### Acute hippocampal slice preparations

Transverse hippocampal slices were obtained from P28 male and female mice (C57BL6/J). Mice were anesthetized with isoflurane and decapitated. The brain was rapidly removed and placed in ice-cold dissection buffer containing (in mM) 252 sucrose, 3 KCl, 7 MgCl_2_, 0.5 CaCl_2_, 1.25 NaH_2_PO_4_, 26 NaHCO_3_, and 25 glucose. The cerebellum was removed with a 20°–30° angle between dorso-ventral and caudo-rostral directions and the resulting flat cut surface was glued onto a precooled vibratome plate; 350 µm sections were collected in the ice-cold dissection buffer using a Leica vibrating microtome VT1200S and maintained at 34°C for 1–2 h in ACSF containing (in mM) 125 NaCl, 2.5 KCl, 26 NaHCO_3_, 1.25 NaH_2_PO_4_, 2 CaCl_2_, 1 MgCl_2_, and 25 glucose. Dissection buffer and ACSF were bubbled with 95% O_2_/5% CO_2_.

### Electrophysiological Recordings

Whole-cell patch-clamp recordings were carried out from CA1 neurons in acute hippocampal slices placed on the stage of a Nikon Eclipse FN1 upright microscope at room temperature and using a Multiclamp 700B amplifier (Axon Instruments). Micropipettes with a resistance in the range of 4–6 MΩ were pulled from borosilicate glass capillaries (Harvard Apparatus 30-0062, GC150T-10) using a micropipette puller (Narishige PC-10 model). The recording chamber was continuously perfused with ACSF bubbled with 95% O2/5% CO2 containing (in mM) 125 NaCl, 2.5 KCl, 26 NaHCO_3_, 1.25 NaH_2_PO_4_, 2 CaCl_2_, 1 MgCl_2_, and 25 glucose. The internal solution (285 mOsm with pH 7.25) contained 125 mM CsMeSO_4_, 10 mM CsCl, 2.5 mM MgCl_2_, 0.4 mM Na-GTP, 4 mM Na_2_ATP, 0.6 mM EGTA, 10 mM Hepes. CA1 pyramidal neurons were identified with DIC. EPSCs were evoked using a bipolar electrode in borosilicate theta glass filled with ACSF and placed in the stratum radiatum. When recording EPSCs, 20 µM bicuculline was added to block inhibitory synaptic transmission. AMPA receptor-mediated currents were recorded at −70 mV and NMDA receptor-mediated currents were recorded at +40 mV and measured 50 ms after the stimulus. The series resistance Rs was left uncompensated. Recordings with Rs higher than 30 MΩ were discarded. PPR was determined by delivering two pulses separated by 50 ms. PPR was defined as the peak current of the second EPSC over the peak current of the first EPSC. EPSCs amplitude and PPR measurements were performed using Clampfit (Axon Instruments).

### Dissociated hippocampal cultures, electroporation, and infection

Primary cultures were prepared from hippocampi of P0 mice as described previously (71), with some modifications. Briefly, hippocampi were dissected out in Hibernate medium and incubated with papain (Sigma-Aldrich, #S3125) for 20 min at 37°C, then dissociated in Hibernate A with 1.8 mM CaCl_2_. Dissociated neurons were plated at a density around 500,000 cells per 60-mm dish on poly-L-lysine and laminin (1 mg.mL^-1^, Sigma-Aldrich, #P2636, #11243217001) precoated glass coverslips (18 mm 1.5H, Marienfield, #117580), into Neurobasal A medium (NbA, Gibco, #12349-015) supplemented with SM1 (StemCell, #05711), 0.5 mM GlutaMAX, and 10% horse-serum (Gibco, #35050-038, #26050-088). After 30 min, or 90 min when neurons were electroporated, coverslips were flipped onto 60-mm dishes containing an astrocyte feeder layer, in NbA supplemented with SM1 and 0.5 mM GlutaMAX, and cultures were maintained at 37°C in 5% CO_2_. Ara-C (2 µM, Sigma-Aldrich, #C1768) was added at DIV-3 to inhibit glial proliferation. Astrocyte feeder layers were prepared 2 weeks in advance from P0 WT mice, plated at a density of 75,000 cells per 60 mm dish previously coated with 0.1 mg/mL^-1^ poly- L-lysine, and cultured at 37°C with 5% CO_2_ in minimal essential medium (MEM, Gibco, #11380037) supplemented with glucose (4.5 g/L), 2 mM GlutaMAX, and 10% horse serum. For biochemistry experiments, hippocampal neurons were plated at a density of 700,000 cells per well in 6-well plates pre-coated with 1 mg/mL^-1^ poly-L-lysine, into Neurobasal A medium (NbA) supplemented with SM1, 0.5 mM GlutaMAX, and 10% horse-serum for the first 30 min, then maintained with the same medium containing only 3% horse-serum at 37°C with 5% CO_2_. After 3-7 days in vitro (depending on the batch of SM1 supplement) the culture medium was replaced with NbA without horse serum to slow glial proliferation. If necessary, Ara-C (2 µM) was added at DIV 6-8 to completely stop glial cells proliferation.

For Figs. 5, 6, S7, S8 and S9, neurons were electroporated prior to plating on coverslips with the Amaxa system (Lonza) using 500,000 cells per cuvette. Depending on the experiments, the following plasmid combinations were used: 1/ BirA^ER^ + Xph20-mRuby2 + GPHN.FingR-GFP (1:1:1 µg DNA); 2/ BirA^ER^ + Xph20-GFP or BirA^ER^ + GPHN.FingR-GFP (2:1 µg DNA); 3/ BirA^ER^ + shNLGN1-GFP or BirA^ER^ + shP53-GFP (2:1 µg DNA). For Figs. 4 and S6, neuron cultures were infected with BirA^ER^-HA- IRES-GFP or IRES-GFP AAV1 at a multiplicity of infection (MOI) of 30,000 (DIV 2 to 6), directly in 6-well plates for mixed cultures, or by incubating coverslips overnight in 12-well plates with 0.5 mL of pre- conditioned Neurobasal A complete medium containing viruses for banker cultures. Coverslips were then returned to the 60-mm dishes. Dissociated cultures transduced with BirA^ER^ either by electroporation or infection were maintained in medium supplemented with 10 µM D-biotin (Sigma-Aldrich, #B4639).

### Neuronal lysates and brain tissue fractionation for biochemistry

Fresh cortex were dissected out and immediately frozen at -80°C. On the day of the experiment, cortices were thawed on ice and ground for 20 s with the FastPrep-24 system (MP Biomedicals) in lysis buffer (50 mM Tris-HCl pH 7.5, 150 mM NaCl, 10 mM EDTA, 1% NP-40, 1% SDS, 0.5% DOC) containing protease inhibitor Cocktail Set III (Merck Millipore, #539134).

Neuronal cultures infected with AAVs in 6-well plates were rinsed in ice-cold PBS and scraped into 70 µL of RIPA buffer (50 mM HEPES pH 7.2, 10 mM EDTA, 1% NP-40, 0.1% SDS, 0.5% DOC) containing protease inhibitors. For co-pull-down experiments, a mild, non-denaturating buffer was used (50 mM Tris-HCl pH 7.5, 150 mM NaCl, 0.5% Triton-X100).

Neuronal cultures plated for surface biotinylation experiments were cooled on ice for 10 min, rinsed in ice-cold DPBS 0 CaCl_2_ 0 MgCl_2_ (Gibco, #14200059) supplemented with 5 mM glucose, and treated with 1 mL of Sulfo-NHS-LC-Biotin EZ-Link (0.33 mg/mL, Thermo Scientific, #21335) for 10 min at 4°C. Cultures were then rinsed in ice-cold DPBS supplemented with 100 mM glycine and scraped in 70 µL of lysis buffer (50 mM Tris-HCl pH 7.5, 150 mM NaCl, 1% Triton-X100, 0.1% SDS) containing protease inhibitors.

All homogenates were kept for 30 min on ice, then centrifuged at 8,000 x g for 15 min at 4°C to remove cell debris. Protein concentration was estimated using the Bradford method (Bio-Rad Protein Assay Kit II, #5000002), or the Direct Detect Infrared Spectrophotometer (Merck Millipore), and protein amounts for inputs were adjusted to prepare 10-20 µg protein samples.

### Protein pull-down by streptavidin beads

For the pull-down of biotinylated bAP-NLGN1, 500 µg of total protein were incubated with 20 µL of streptavidin magnetic beads (Pierce, #88817). After 1 h of incubation (rotating wheel, RT), tubes were placed in a magnetic column and beads were washed three times with lysis buffer. Proteins were eluted from the beads by adding 25 μL of Laemli Sample Buffer 2X (Sigma-Aldrich, #S3401), before heating tubes at 95°C. Samples were then processed for SDS-PAGE and Western blotting. To isolate surface-labeled biotinylated proteins, 450 µg of total protein were incubated for 20 min (rotating wheel, RT) with 50 µL of streptavidin-agarose beads (Sigma-Aldrich, #S1638). After incubation, beads were washed three times with lysis buffer, then centrifuged (1,000 x g for 1 min at 4°C) to isolate them. The same beads were used to detect the synaptic partners of bAP-NLGN1 by co-PD, by incubating 900 µg of total protein with 45 µL of beads for 30 min (rotating wheel, 4°C). To saturate the beads in the negative condition (BirA-), 3 µg of free biotin (Sigma-Aldrich, #B4639) was added to the lysate during the incubation. The same amount of biotin was added in the positive condition (BirA+) for 5 min during the washes. Beads were washed 3 times with wash buffer (PBS containing 0,02% Tween20) and isolated. In cellulose acetate filter spin cups (Pierce, #69702), proteins were eluted from the agarose beads by adding 25 μL of Laemli Sample Buffer 2X, heating at 95°C, and finally centrifuging the spin cups. Samples were then processed for SDS-PAGE and Western blotting.

### SDS-PAGE and Western blot

Samples were loaded on 4-20% gradient gels (PROTEAN TGX Precast Protein Gels, BioRad) for separation (200 V, 400 mA, 40 min) and then transferred to nitrocellulose membranes for semi-dry immunoblotting (7 min, TurboBlot system, BioRad). For total protein quantification, membranes were rinsed in water, incubated with the REVERT Total Protein Stain reagents (LI-COR, #926-11010), and imaged immediately in the 700 nm channel with an Odyssey Fc imaging system (LI-COR). Then, membranes were rinsed in Tris-buffered saline Tween-20 (TBST; 28mM Tris, 137mM NaCl, 0.05% Tween-20, pH 7.4) and blocked with 5% non-fat dried milk for 45 min at RT. After blocking, membranes were incubated for 1 h at RT, or overnight at 4°C, with the primary antibody diluted in TBST solution containing 0,5% dry milk: rabbit anti-NLGN1 (Synaptic Systems, #129-013, 1:1000); rabbit anti-NLGN2 (Synaptic Systems, #129-202, 1:1000) ; rabbit anti-NLGN3 (Synaptic Systems, #129-113, 1:1000); rabbit anti-MDGA1 (Synaptic Systems, #421-002, 1:500); mouse anti-PSD-95 (Invitrogen, clone 7E3-1B8 #MA1-046, 1:1000); mouse anti-gephyrin (Synaptic Systems, clone 3B11 #147-111, 1:2000); mouse anti-GluA1 (Neuromab, #75-327, 1:1000); mouse anti-GluA2 (Sigma-Aldrich, #MAB397, 1:1000) ; mouse anti-GluN1 (Sigma-Aldrich, clone 54.1 #MAB363, 1:1000); mouse anti-βactin (Sigma-Aldrich, #A5316, 1:5000); mouse anti-HA (BioLegend, #901513, 1:1000); mouse anti-GFP (Roche, #11814460001, 1:1000); rabbit anti-PSD-95 (Cell Signaling, clone D27E11 #3450, 1:1000). After three washes in TBST, membranes were incubated with horseradish peroxidase (HRP)-conjugated donkey anti-mouse or anti-rabbit secondary antibodies (Jackson Immunoresearch, #715-035-150 and #711- 035-152, 1:5000) for 1 hr at RT. Target proteins were detected by chemiluminescence with SuperSignal West Femto (Pierce) or Clarity Western ECL Substrate (Bio-Rad) on the Odyssey Fc imaging system (LI-COR). The intensity of input signals was normalized to β-actin or total protein, whereas PD signals were normalized on streptavidin beads quantity (total protein staining).

### Immunocytochemistry

To visualize pre- and post-synaptic markers in WT or KI mice cultures, neurons were rinsed in Tyrode solution (15 mM D-glucose, 108 mM NaCl, 5 mM KCl, 2 mM MgCl_2_, 2 mM CaCl_2_ and 25 mM HEPES, pH = 7.4, 280 mOsm) before being fixed for 10 min in 4% paraformaldehyde, 4% sucrose, quenched for 15 min in NH_4_Cl 50 mM in PBS, and permeabilized for 5 min with 0.1% Triton X-100 in PBS. After blocking during 30 min in PBS containing 0.5% BSA, 0.1% Triton X-100, and 2% Goat Serum (Gibco, #PCN5000), neurons were stained for 90 min with the following primary antibodies: mouse anti-PSD- 95 (Invitrogen, #MA1-046, 1:400) and guinea pig anti-VGLUT1 (Merck Millipore, #AB5905, 1:2000), or mouse anti-gephyrin (Synaptic Systems, #147-111, 1:400) and guinea pig anti-VGAT (Synaptic Systems, #131-004, 1:1000). For each combination, the neuronal microtubule cytoskeleton was labeled using rabbit anti-MAP-2 (Merck Millipore, #AB5622, 1:800). Following three washes, neurons were incubated for 45 min with appropriate secondary antibodies: goat anti-mouse DL405 (Jackson ImmunoResearch, #115-475-003, 1:800), goat anti-guinea pig AF568 (Invitrogen, #A11075, 1:800), and goat anti-rabbit AF488 (Invitrogen, #A11008, 1:400).

To visualize endogenous PSD-95 and gephyrin in comparison with the intrabodies, neurons were treated as above and stained with the following antibodies: mouse anti-PSD-95 (Invitrogen, #MA1- 046, 1:400) or mouse anti-gephyrin (Synaptic Systems, #147-111, 1:400), combined with rabbit anti-MAP-2 (Merck Millipore, #AB5622, 1:800), followed by goat anti-mouse Dylight405 (Jackson ImmunoResearch, #115-475-003, 1:800) and goat anti-rabbit AF568 (Invitrogen #A11011, 1:800). Coverslips were mounted in Fluoromount-G (SouthernBiotech, #0100-01) and immunostainings in primary cultures were visualized on an inverted microscope (Nikon Eclipse TiE) equipped with a 60x/1.40 NA objective and filter sets for Dylight405 (Excitation: FF02-379/34; Dichroic: FF-409Di03; Emission: FF01-440/40); Alexa Fluor 488 (Excitation: FF01-472/30; Dichroic: FF-495Di02; Emission: FF01-525/30); and Alexa Fluor 568 (Excitation: FF01-543/22; Dichroic: FF-562Di02; Emission: FF01- 593/40) (SemROCK). Images were acquired with a sCMOS camera (PRIME 95B, Photometrics) driven by the Metamorph software (Molecular Devices). The density and surface area of PSD-95, VGLUT1, gephyrin, and VGAT puncta was measured using a custom macro written in Metamorph, as described (72).

### Labeling of biotinylated bAP-NLGN1 with fluorescent streptavidin

To visualize endogenous bAP-NLGN1 proteins at the membrane, neurons were rinsed in Tyrode solution and blocked with 1% biotin-free Bovin Serum Albumin (BSA, Roth, #0163) for 10 min at 37°C. Neuron coverslips were then incubated live with AF647-conjugated streptavidin (Invitrogen, #S32357, 4 µg/mL) for 10 min at 37°C, rinsed, and either mounted for observation in an open Inox observation chamber (Ludin, Life Imaging Services) containing pre-warmed Tyrode solution, or processed for dSTORM imaging. Live-labeling in primary cultures were visualized on an inverted microscope (Nikon Eclipse TiE) equipped with an APO TIRF 100x/1.49 NA oil immersion objective and enclosed in a thermostatic box (Life Imaging Services) providing air at 37°C. Fluorescence signals were detected using a mercury lamp (Nikon Xcite) and filter sets for EGFP (Excitation: FF01-472/30; Dichroic: FF- 495Di02; Emission: FF01-525/30), mRuby2 (Excitation: FF01-543/22; Dichroic: FF-562Di02; Emission: FF01-593/40), and AF647 (Excitation: FF02-628/40; Dichroic: FF-660Di02; Emission: FF01-692/40) (SemROCK). Images were acquired with an EMCCD camera (Evolve, Roper Scientific) driven by the Metamorph software (Molecular Devices). The quantification of SA-AF647 signal intensity and its colocalization with PSD-95 and gephyrin puncta were measured using a custom macro written in Metamorph.

### Direct Stochastic Optical Reconstruction Microscopy (dSTORM)

Neurons previously labeled with SA-AF647 were fixed for 10 min in 4% paraformaldehyde, 4% sucrose, quenched for 15 min in NH_4_Cl 50 mM in PBS, and kept at 4°C until dSTORM acquisitions. The coverslips were mounted in a Ludin chamber in an oxygen-scavenging imaging buffer: Tris-HCl buffer (pH 7.5), containing 10% glycerol, 10% glucose, 0.5 mg.mL^-1^ glucose oxydase (Sigma-Aldrich, #G2133), 40 mg.mL^-1^ catalase (Sigma-Aldrich, #C100, 0,1% w/v) and 50 mM β-mercaptoethylamine (MEA, Sigma-Aldrich, #M6500) (73) and sealed using a second glass coverslip. The same microscope described above for live-labeling was used, which is further equipped with a perfect focus system preventing drift in the z-axis. A four-color laser bench (405/488/561 nm lines, 100 mW each, Roper Scientific; and 647 nm line, MPB Communications Inc.) is connected through an optical fiber to the Total Internal Reflection Fluorescence (TIRF) illumination arm of the microscope and laser powers are controlled through an Acousto-Optical Tunable Filter (AOTF) driven by Metamorph. The fluorophores AF647 were excited with the 647 nm laser line through a four-band beam splitter (BS R405/488/561/635, SemROCK). Pumping of AF647 dyes into their triplet state was performed for several seconds using ∼60 mW of the 647 nm laser at the objective front lens. Then, a lower power (∼20 mW) was applied to detect the stochastic emission of single-molecule fluorescence, which was collected using the optics and detector described above and a FF01-676/29 nm emission filter (SemROCK). 10 streams of 4,000 frames each were acquired at 50 Hz (20 ms exposure time) using Metamorph. Nano-diamonds (Adamas Nanotechnologies, #ND-NV140 nm) were added to the samples for later registration of images and lateral drift correction. The localization precision of our imaging system in dSTORM conditions is around 60 nm (FWHM). Analysis of the image stacks was made offline under Metamorph, using the PALM-Tracer program based on wavelet segmentation for single-molecule localization. Unique super-resolved image of 32 nm pixel size (zoom 5 compared to the original images) was reconstructed by summing the intensities of all localized single molecules (1 detection per frame is coded by an intensity value of 1). The analysis of protein enrichment at PSD-95 or gephyrin positive synapses was performed using a custom macro written in Metamorph, allowing the count of the average number of detections within synaptic puncta in comparison with the average number of extra-synaptic detections. The SR-Tesseler software (49) was used to identify bAP-NLGN1 domains into regions of interest (PSD-95 or gephyrin puncta). This method uses a Voronoï diagram to decompose a super-resolution image into polygons of various sizes centered on the localized molecules.

### Organotypic slice culture

Organotypic hippocampal slice cultures were prepared from either WT or KI mice, as described (50). Animals at postnatal days 5–6 were quickly decapitated and brains placed in ice-cold Gey’s balanced salt solution under sterile conditions. Hippocampus were dissected out and coronal slices (350 µm) were cut using a tissue chopper (McIlwain) and incubated at 35°C in 5% CO2 with serum-containing medium on Millicell culture inserts (CM, Millipore). BirA+ AAVs were added at DIV 3 and the medium was replaced every 2–3 days until slices were used at DIV14.

### Labelling of biotinylated bAP-NLGN1 **in** organotypic slices with fluorescent or nanogold-conjugated streptavidin

Organotypic slices of WT or KI mice infected with BirA+ AAVs were live-labeled for 20 min at 35°C with either SA-AF647 (stock 2 mg/mL, 1:400) or StreptAvidin conjugated to 1.4 nm NanoGold particles (SA-Gold, Nanoprobes Inc., stock 80 µg/mL, 1:100) in ACSF solution containing 1% biotin-free BSA. Slices were fixed with 4% PFA-sucrose and 0.2% glutaraldehyde in 1X PBS overnight at 4°C. Slices treated with SA-AF647 were observed by confocal microscopy, as described above for immunolabeled brain slices, while samples treated with SA-Gold were subjected to silver enhancement for 10 min (LI Silver, Nanoprobes Inc). Samples were then post-fixed with 1% glutaraldehyde in 0.15 M Sorensen’s phosphate buffer (SPB; Electron Microscopy Sciences) for 10 min at room temperature, then incubated with 1.5% OsO_4_ and 1.5% potassium ferrocyanide for 30-min on ice. Sequential dehydration was performed with 70%, 90%, 95%, and 100% ethanol, followed by one incubation with 100% acetone, then samples were embedded for 2 h with 1:1 acetone and epon resin (Embed-812, EMS), followed by 100% epon resin at 60°C for 48 h. TEM imaging was performed in high contrast mode with a H7650 transmission electron microscope (Hitachi) equipped with an Orius SC1000 CCD camera (Gatan Inc). The number of silver particles within each synapse was manually quantified on Image-J by setting a detection threshold for all images, after subtracting the background.

### Statistical analysis

Statistical significance tests were performed using GraphPad Prism software 9.1.0 (San Diego, CA). For two-condition data sets, comparisons were made using the non-parametric Mann-Whitney test. Kruskal-Wallis test was used to compare more than two groups showing non-normal distributions, and was followed by a post-hoc multiple comparison Dunn’s test. Statistical significance was assumed when p < 0.05, and is represented as: *p < 0.05, **p < 0.01, ***p < 0.001, ****p < 0.0001. The number of experiments performed and samples examined are indicated in the relevant figure legends. When the differences between groups were not significant, no symbol was indicated on the graphs.

## Acknowledgements

We thank D. Arnold, M. Sainlos, P. Scheiffele, and A. Ting for the generous gift of DNA plasmids; M. Munier and S. Benquet for molecular biology; C. Breillat, S. Daburon, and D. Choquet for the gift of viral BirA-IRES-GFP and control constructs, J.B. Sibarita, A. Kechkar and C. Butler for the gift of the PALM-Tracer and WAVE-Tracer programs; M.C. Birling at the Phenomin facility (Strasbourg) for the generation of the bAP-NLGN1 KI mouse strain; P. Costet, C. Martin, and H. El Oussini at the Plateforme In Vivo Exempte d’Organisme Pathogène Spécifique (PIV-EOPS) and the “Plateforme In Vivo experimental” (PIV-EXPE) of the Bordeaux Neurocampus for maintenance of the mouse strain and completion of the SHIRPA tests; D. Gonzales at the genotyping facility of Bordeaux Neurocampus (NeuroCentre Magendie); E. Verdier, N. Retailleau, A. Caralp, and L. Leroy at the Cell Biology facility of the Bordeaux Neurocampus (IINS), N. Dutheil at the Virology platform of the Bordeaux Neurocampus (IMN) for the production of AAVs; M. Petrel, S. Lacomme, and M. Fernandez-Monreal (Bordeaux Imaging Center) for advice on TEM; M. Mondin (Bordeaux Imaging Center) for Metamorph macros; R. Sterling, A. Castets and Z. Andrieux for logistics (IINS); M. Sainlos, S. Levi, and M. Montcouquiol for insightful discussions.

This work received funding from the Centre National de la Recherche Scientifique (CNRS), Agence Nationale pour la Recherche (grants “Synaptoligation” and “NanoSynAtlas”), Conseil Régional Aquitaine, Fondation pour la Recherche Médicale (“Equipe FRM” DEQ20160334916 and Postdoctoral fellowship to C. Ducrot #SPF202209015897), national infrastructure France BioImaging (grant ANR- 10INBS-04-01), Labex BRAIN (ANR-10-LABX-43), and GPR BRAIN_2030.

## Author contributions

C.D^1^. performed tissue collection, immunohistochemistry, confocal and electron microscopy and performed some behavioral experiments; A.D. performed biochemical pull-down, immunocytochemistry, and super-resolution imaging; C.D^2^. and B.T. prepared primary hippocampal cultures, organotypic slices, and performed biochemistry; C.D^1,2^, A.D., and B.T. performed viral infection of dissociated and organotypic hippocampal cultures; M.L. performed electrophysiology; O.T., A.F., and M.L. provided funding. O.T. coordinated the research project, performed preliminary experiments, and prepared the manuscript draft. All authors discussed the results and participated to the manuscript writing.

Author information : ^1^Charles Ducrot; ^2^Chloé Desquines.

## Competing financial interests

The authors declare no competing financial interest.

## Notes

### Competing Interest Statement

The authors have declared no competing interest.

