## Supplemental figures for "High-affinity detection of endogenously biotinylated neuroligin-1 at excitatory and inhibitory synapses using a tagged knock-in mouse strain"

**Affiliations:**
1. Univ. Bordeaux, CNRS UMR 5297, Interdisciplinary Institute for Neuroscience, IINS, F-33000 Bordeaux, France

### Supplementary figures

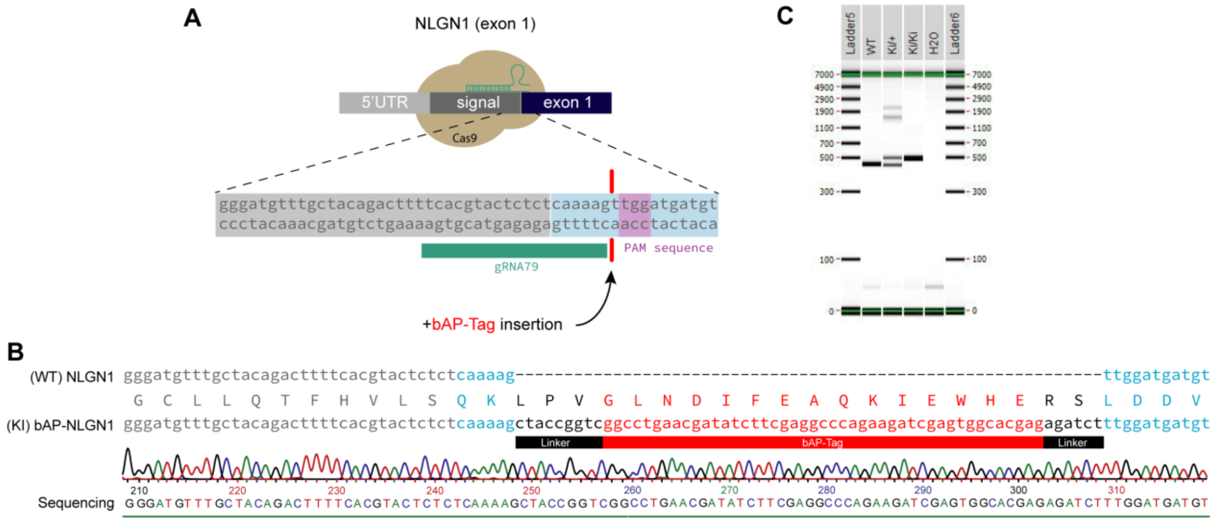


*Figure S1. Generation and validation of the bAP-NLGN1 KI mice*

**(A)** Schematics showing the specific sequence targeted by the gRNA in the signal peptide sequence located in the exon 1 of the Nlgn1 gene. **(B)** Representative genomic DNA sequencing showing the correct insertion of the bAP-tag in the NLGN1 gene. **(C)** Example of PCR genotyping results obtained from homozygous or heterozygous KI mice, in comparison with WT animals.

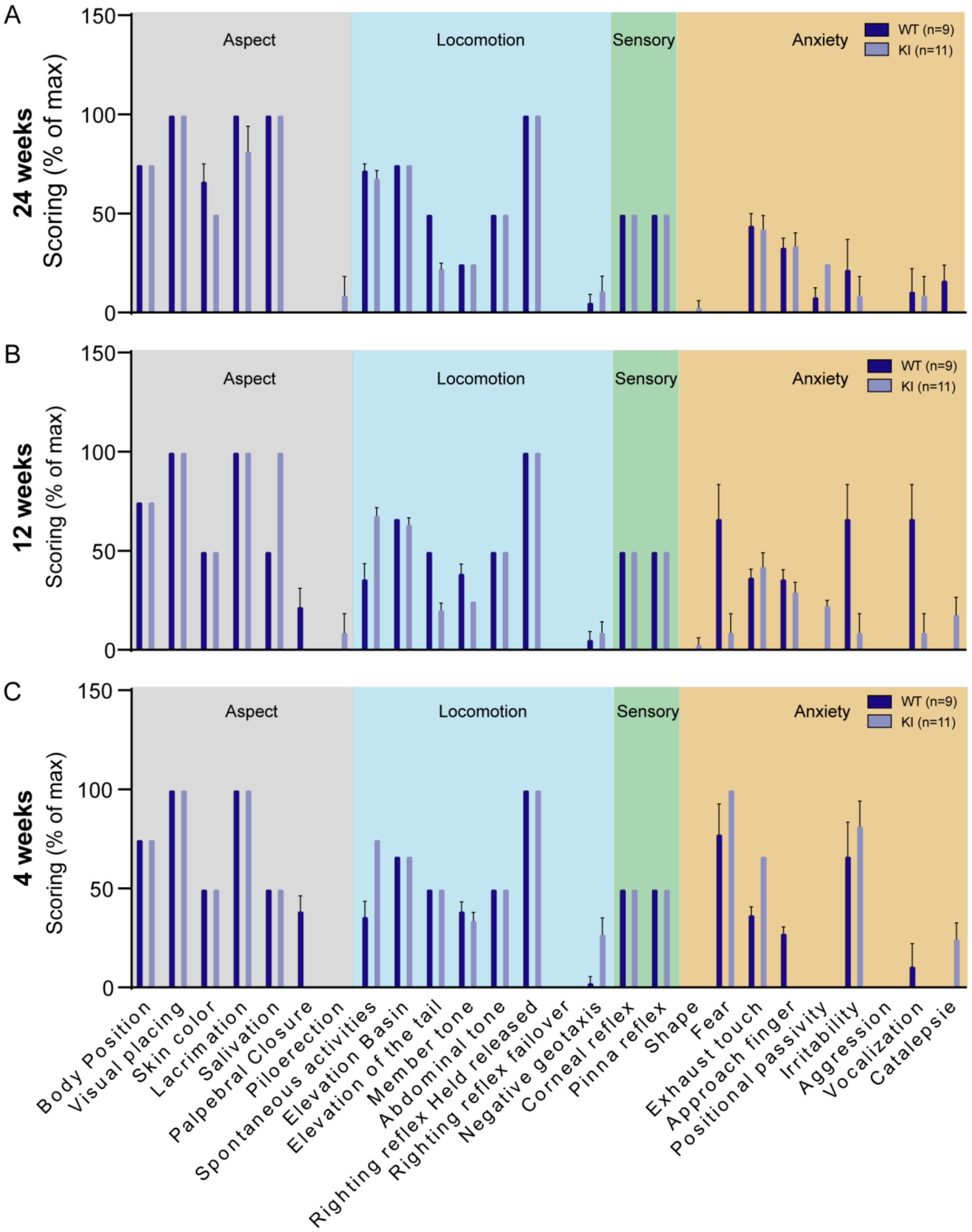


***Figure S2. Detailed analytical results from the SHIRPA tests.***

**(A-C)** Detailed behavioral screening of KI mice in comparison to control mice (WT) at 4, 12, and 24 weeks of age, taking into consideration their general aspect, anxiety, locomotion and sensory abnormalities. n = 9 WT mice and 11 KI mice.

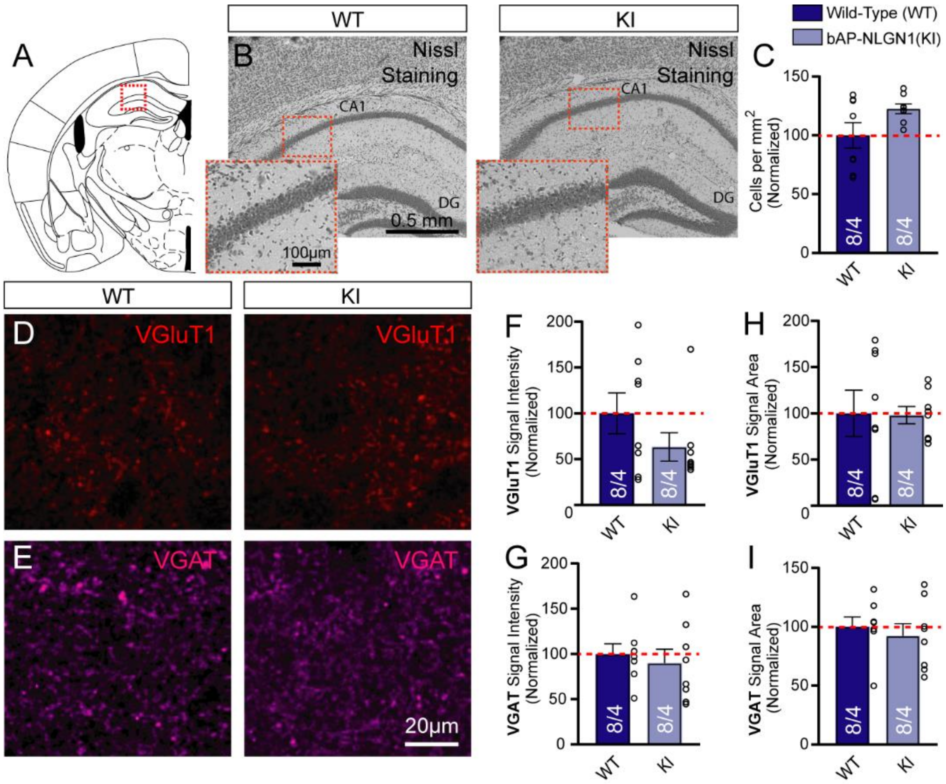


***Figure S3. Immunostaining of excitatory and inhibitory synapses in hippocampal slices from WT and KI mice***

**(A)** Schematic representation of the hippocampal slices used for immunohistochemistry characterization (*bregma -1.9 mm*). Surface and intensity for each signal were measured in a series of 2 different hippocampal slices ranging from Bregma -1.5 to -1.9 mm with a total of 4 areas of each hemisphere. **(B)** Representative photomicrographs of Nissl staining in the CA1 regions of WT and KI brain sections, respectively. **(C)** Number of cells per $mm^2$ in the hippocampus from WT and KI mice brain sections ($p$ = 0.23). **(D, E)** Representative confocal images of immunohistochemical staining for VGluT1 (red) and VGAT (magenta), respectively, performed on hippocampal slices from WT and KI mice. Quantification of the signal intensity **(F, G)** and signal surface area **(H, I)**, for VGluT1 and VGAT staining, respectively, normalized to WT levels (in %) (F, $p$ = 0.44; G, $p$ = 0.50; H, $p$ = 0.72; I, $p$ = 0.44). For all experiments (n= 8 hemispheres/4 mice for each genotype). Data represent the mean ± S.E.M and were compared by a Mann-Whitney test.

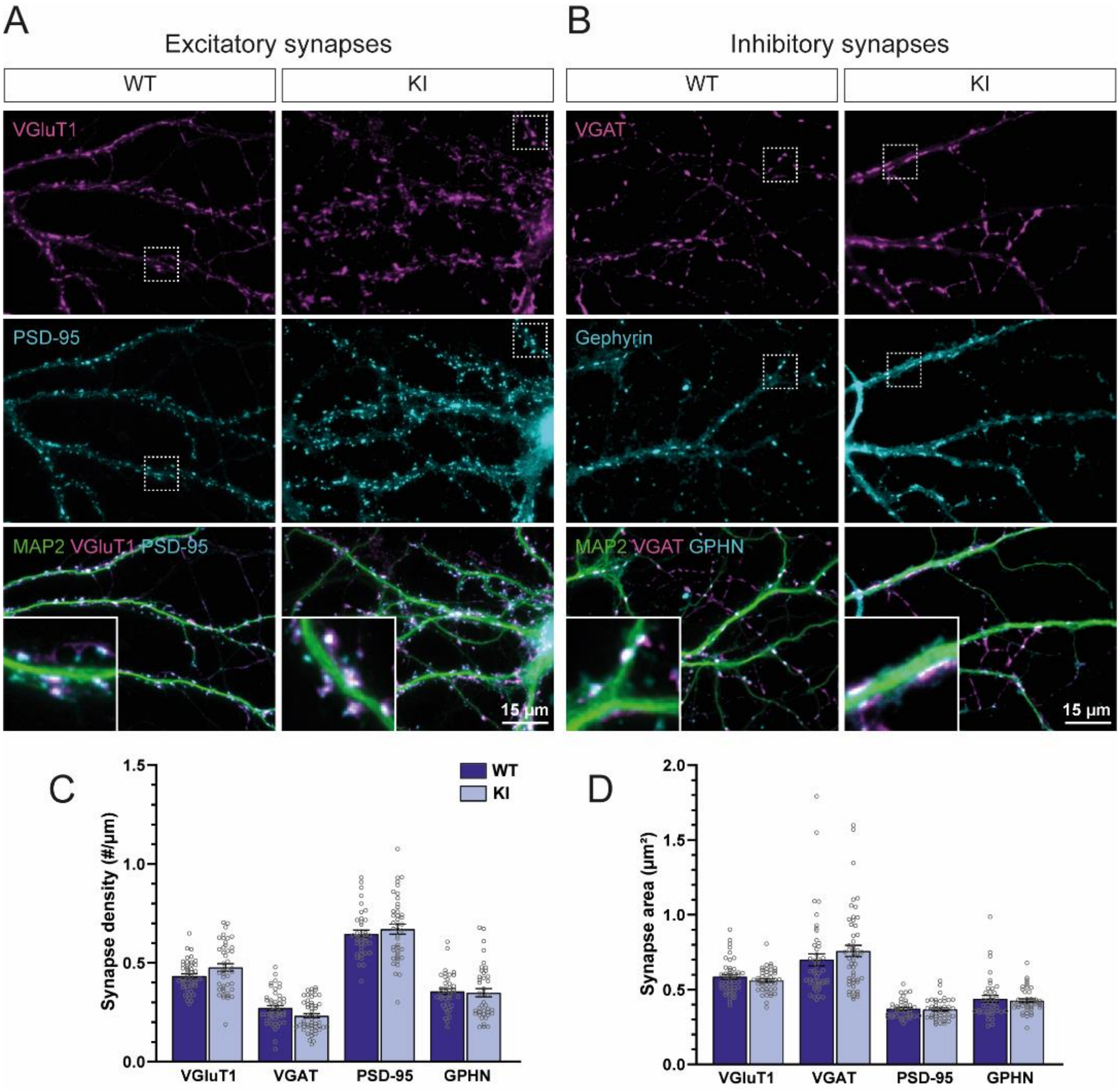


***Figure S4. Synapse development in KI mice cultures***

**(A, B)** Representative images of hippocampal neurons dissociated from WT or KI mice, and processed at DIV 14 for triple immunofluorescence staining of MAP2 to visualize microtubules (green), and of either excitatory or inhibitory synaptic markers, respectively. Antibodies to VGluT1 (magenta) and PSD-95 (cyan) were used to visualize excitatory pre- and post-synapses, respectively, while antibodies to VGAT (magenta) and gephyrin (cyan) were used to visualize inhibitory pre- and post-synapses, respectively. Insets in the merged images show zooms of dendritic segments highlighted by the dashed squares in the lower magnification images. **(C, D)** Bar plots showing the density per unit dendrite length, and surface area respectively, of individual VGluT1, VGAT, PSD-95, and gephyrin puncta, in cultures from WT and KI mice. Dots in the bars represent individual cells, with n > 38 cells for each condition, from 3 independent experiments. Data represent mean ± SEM and were compared by a Kruskal–Wallis test followed by Dunn's multiple comparison test (C, *pVGluT1*, *pPSD-95*, *pGPHN* > 0.99, *pVGAT* = 0.82; and D, *pVGluT1*, *pVGAT*, *pPSD-95*, *pGPHN* > 0.99).

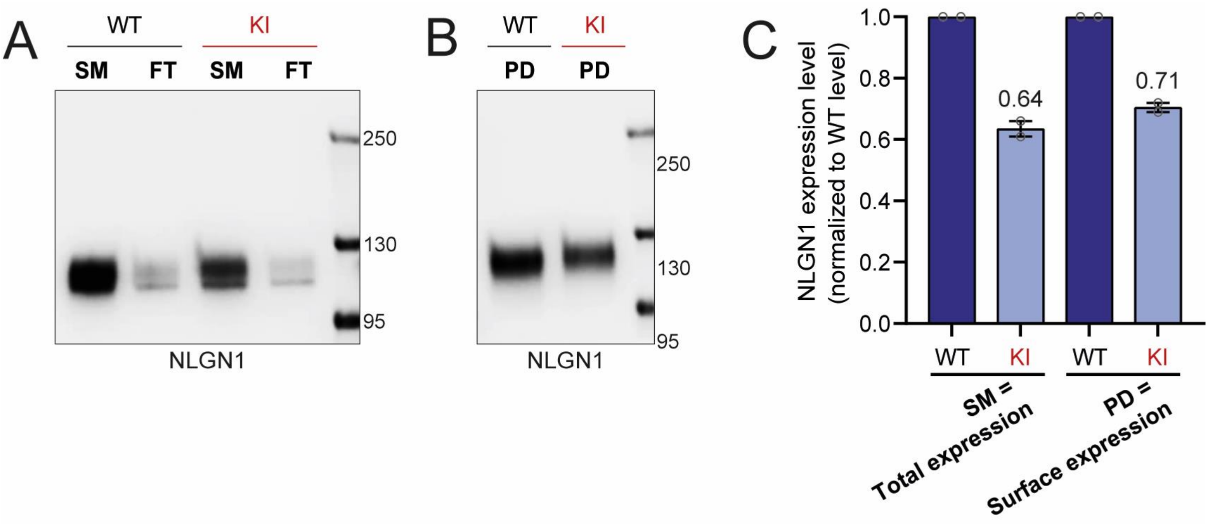


***Figure S5. bAP-NLGN1 expression level at the neuronal surface***

Neuron cultures from WT or KI mice were processed for surface biotinylation at DIV 14, then lysed. Protein extracts were either loaded as starting material (SM) or pulled-down with streptavidin beads (PD). The protein fraction that did not bind to the beads was isolated in the flow-through (FT). **(A, B)** NLGN1 protein was identified by Western blot in both SM and FT fractions, or in the PD fraction, respectively (molecular weight markers in kDa indicated on the right). Total protein staining (not shown) was used as a loading control. **(C)** Relative NLGN1 expression level, in the total protein fraction and in the membrane protein fraction, analyzed by semi-quantitative immunoblotting. Dots in the bars represent individual cultures (n = 2 independent cultures for all experimental groups). Data represent mean ± SEM and were compared by a Kruskal–Wallis test followed by Dunn's multiple comparison test (SM, $p = 0.06$; PD, $p = 0.38$).

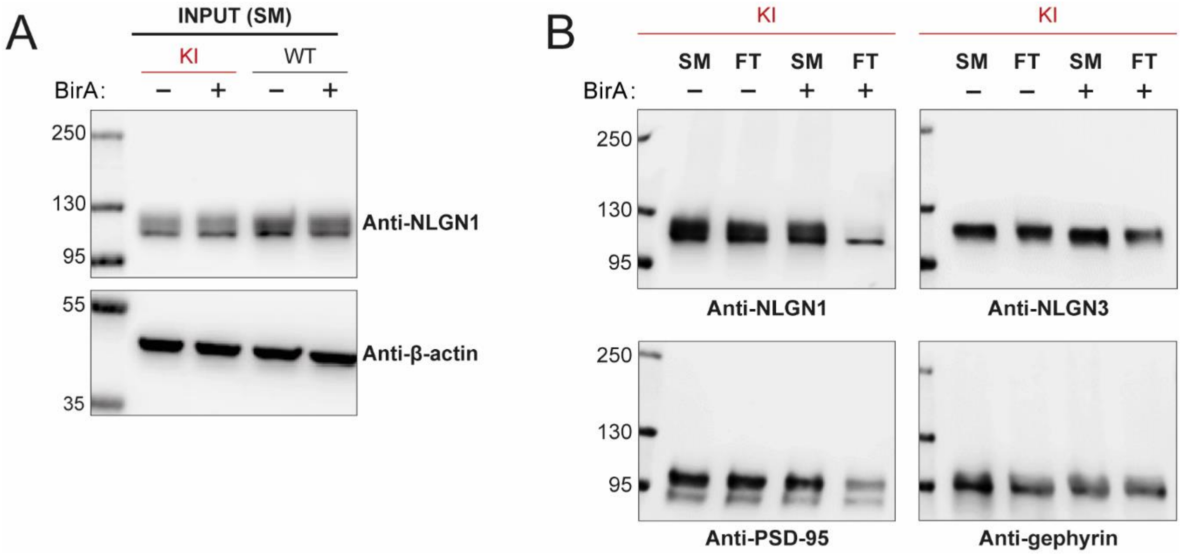


***Figure S6. Inputs from pull-down experiments***

Cultures from WT or KI mice were infected at DIV 3-5 with the BirA$^{ER}$-HA-IRES-GFP virus (BirA+), or the IRES-GFP control virus (BirA-). Neuron cultures were lysed at DIV 14, and protein extracts were either loaded as starting material (SM) or precipitated with streptavidin beads. Protein fraction that did not bind to the beads was isolated in the flow-through (FT). Separated proteins were identified by Western blot (molecular weight markers in kDa indicated on the left). **(A)** NLGN1 protein identified in the SM (top membrane). β-actin was used as loading control (bottom membrane). **(B)** NLGN1, NLGN3, PSD-95 and gephyrin proteins identified in both SM and FT fractions. Total protein staining (not shown) was used as a loading control.

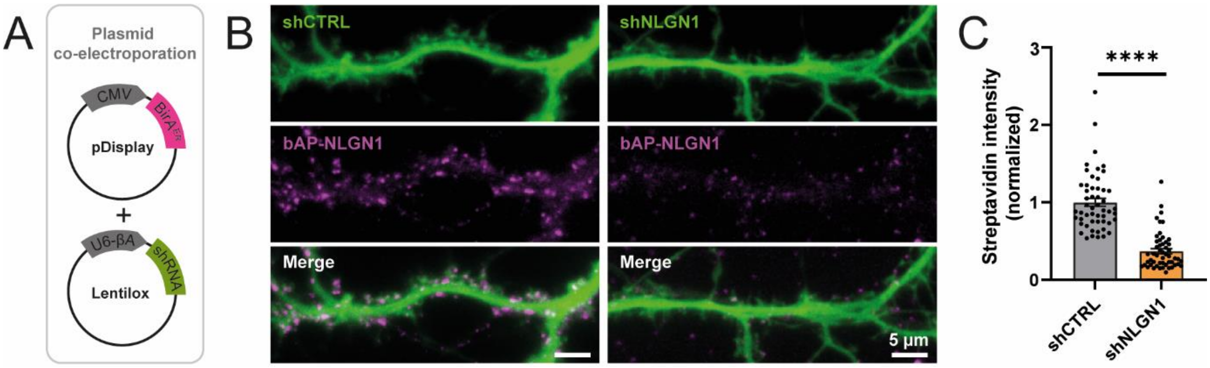


***Figure S7. Knock-down of bAP-NLGN1 reduces streptavidin staining***

**(A)** Dissociated hippocampal neurons from KI mice were electroporated at DIV 0 with a combination of $BirA^{ER}$ and shRNAs to either p53 (shCTRL) or to NLGN1 (shNLGN1), both containing a GFP reporter. **(B)** At DIV 14, neurons were live-labeled with SA-AF647 and visualized by epifluorescence microscopy. Representative images of dendritic segments showing the GFP signal revealing the shRNA expression (green), bAP-NLGN1 labeling with SA-AF647 (magenta), and merged images. **(C)** Quantification of the SA-AF647 fluorescence intensity in each electroporation condition. Dots in the bars represent individual cells, with n = 52 cells for each condition, from 2 independent experiments. Data represent mean ± SEM and were compared by a Mann-Whitney test (****$P < 0.0001$).

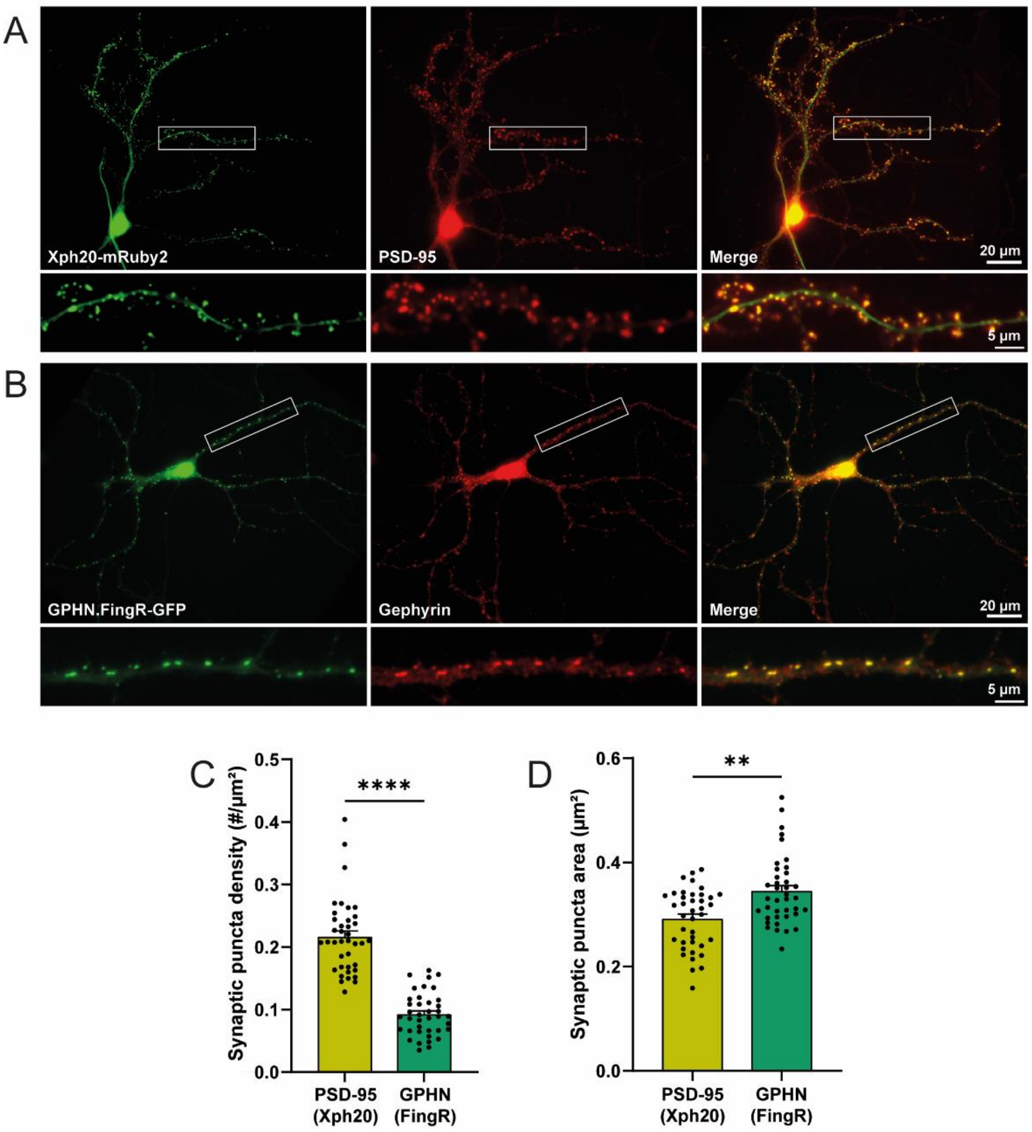


#### *Figure S8. Validation of PSD-95 and gephyrin intrabodies*

Dissociated hippocampal neurons from bAP-NLGN1 mice were electroporated at DIV 0 with Xph20-mRuby2 and GPHN.FingR-GFP encoding plasmids, then immunolabeled with PSD-95 or gephyrin antibodies. **(A, B)** Representative neurons showing either the Xph20-mRuby2 intrabody against PSD-95 (green), and endogenous PSD-95 (red), or the GPHN.FingR-GFP against gephyrin (green), and endogenous gephyrin (red), respectively. Merged images on the right highlight the colocalization of both signals (yellow). Dendritic segments corresponding to the rectangle areas are shown in the bottom images. **(C, D)** Bar plots showing the mean density per unit dendrite area and individual surface area, of Xph20-mRuby2 and GPHN.FingR-GFP synaptic puncta, respectively (dots in the bars represent individual cells, with n = 39 cells for each condition, from 2 independent experiments). Data represent mean ± SEM and were compared by a Mann-Whitney test (**$P < 0.01$, ****$P < 0.0001$).

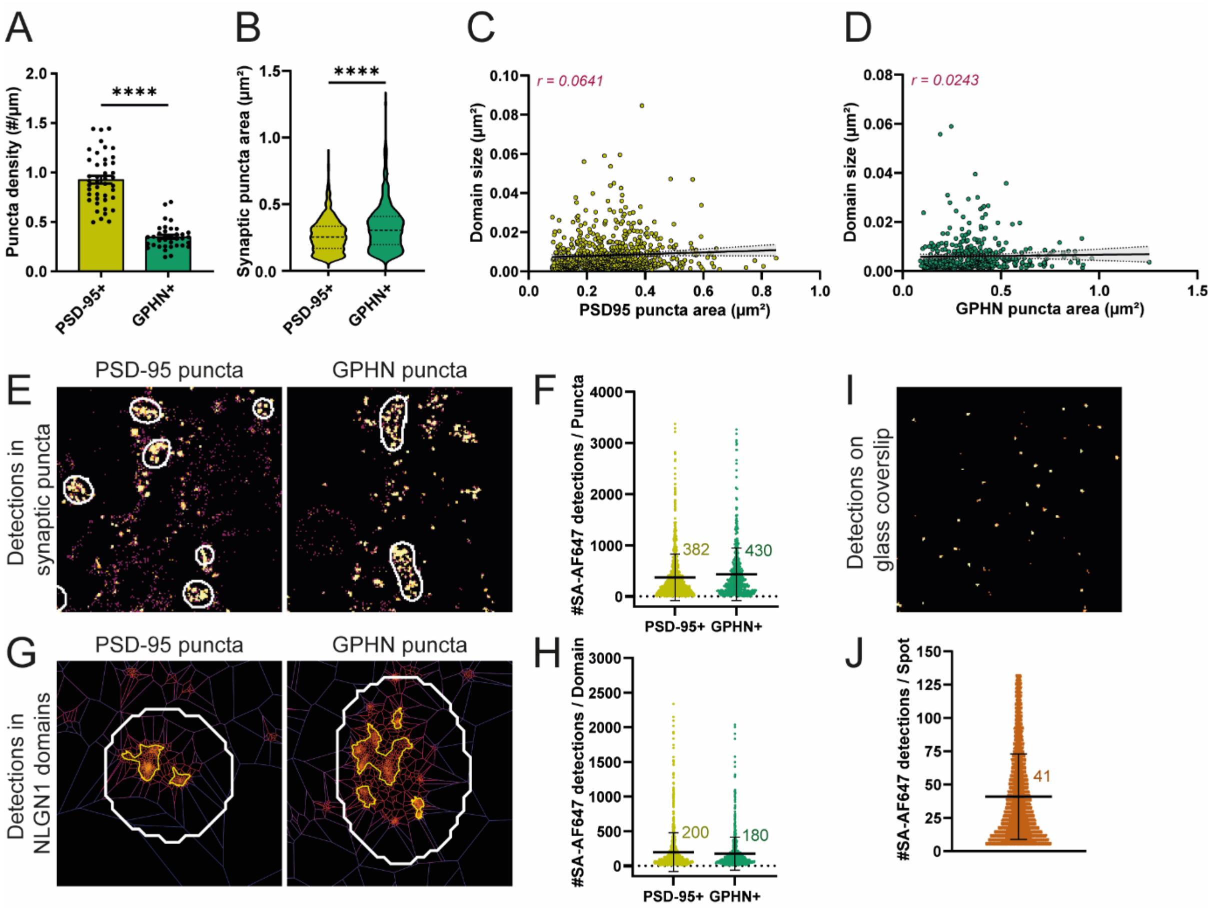


### *Figure S9. Quantitative parameters from dSTORM experiments*

**(A-H)** Dissociated hippocampal neurons from bAP-NLGN1 mice were electroporated at DIV 0 with a combination of BirA$^{ER}$ and either Xph20-GFP or GPHN.FingR-GFP. dSTORM experiments were performed at DIV 14 on fixed cultures, after live-labeling neurons with SA-AF647. **(A)** Quantification of the density per unit dendrite length of individual PSD-95 and gephyrin positive puncta. Data represent mean ± SEM and dots represent individual cells. **(B)** Violin plots showing the surface area of individual PSD-95+ or gephyrin synaptic puncta. Data were compared by a Mann-Whitney test (****P < 0.0001). **(C, D)** Absence of correlation between the bAP-NLGN1 domain area and PSD-95 or gephyrin puncta area, respectively. Dots represent individual synaptic puncta. r : Pearson's correlation coefficient; black line: linear regression with 95% confidence interval (grey). **(E)** Representative images of dendritic segments showing bAP-NLGN1 detections (gold) enriched in PSD-95 or gephyrin positive synapses (outlines in white). **(G)** Examples of bAP-NLGN1 localization organized in a subset of nanodomains (outlines in gold) within PSD-95 and gephyrin positive puncta (outlines in white). **(F, H)** Number of SA-AF647 detections at individual PSD-95 or gephyrin positive post-synapses, or at individual bAP-NLGN1 nanodomains within these post-synapses, respectively. Dots represent individual synaptic puncta or individual nanodomains, respectively. All data related to PSD-95+ puncta were obtained from 4 independent experiments, with 1390 domains / 1065 synaptic puncta / 43 cells analyzed. Data related to GPHN+ puncta were obtained from 3 independent experiments, with 984 domains / 631 synaptic puncta / 36 cells analyzed. **(I, J)** dSTORM experiments were performed on highly diluted SA-AF647 immobilized on glass coverslips. **(I)** Representative image of SA-AF647 detections (gold), each spot indicating a SA-AF647 molecule detected several times. **(J)** Number of SA-AF647 detections at individual spots (n = 17237 spots, from 3 glass coverslips). Data in **F**, **H**, **J** represent mean ± SD (mean values written on the side).

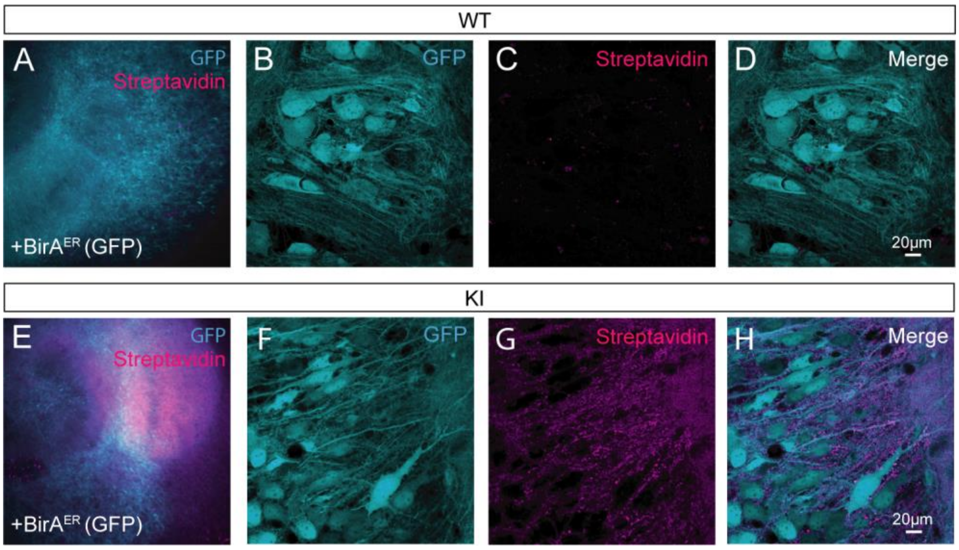


***Figure S10. Visualization of bAP-NLGN1 in organotypic brain slices***

**(A, E)** Representative images (10X) of the CA3 region in hippocampal slice cultures from WT **(A)** or KI **(E)** mice infected with BirA$^{ER}$-IRES-GFP AAV9 (cyan) and labeled with SA-AF647 (magenta). **(B-D** and **F-H)** Representative confocal images (63X) of CA3 pyramidal neurons in organotypic slice cultures from WT or KI mice, respectively, infected with BirA$^{ER}$-IRES-GFP AAV9 (cyan) (**B, F**) and labeled with SA-AF647 (magenta) (**C, G**). Images in **(D, H)** show the merged colors.
